# Urinary proteins from Sickle Cell patients induce inflammation and kidney injury via the TGFβ-p53 axis in a podocyte cell culture model

**DOI:** 10.1101/2025.08.08.669286

**Authors:** Wasco Wruck, Abida Islam Pranty, Chantelle Thimm, Christiane Loerch, Rosanne Mack, Theresa Koranteng-Adjaye, Vincent Boima, Yvonne Dei-Adomakoh, Margaret Lartey, James Adjaye

## Abstract

**Background:** Sickle cell disease (SCD) is an inherited blood disorder affecting the oxygen-carrying hemoglobin in red blood cells making them deform into a sickle shape. Hemolysis and vaso-occlusion associated with this process can lead to complications in many organs and frequently to renal complications. Numerous factors are considered to contribute towards the development of proteinuria (PU) in SCD including hyperfiltration, ischemia, oxidative stress and decreased nitric oxide (NO) bioavailability but the detailed pathophysiology still needs further elucidation.

**Methods:** Employing arrays, we investigated cytokines and kidney injury-associated markers in the urine of a cohort of SCD patients from Ghana carrying the SS and SC genotypes which were further sub-divided into groups with proteinuria (SCD_PU) and without proteinuria (SCD).

**Results:** We identified up-and down-regulated proteins when comparing SCD with and without proteinuria. Amongst these is the well-established kidney injury marker-Clusterin which was up-regulated and could be validated in an ELISA-based assay. Refining the study to the SS and SC genotypes, we identified (and confirmed by ELISA) another established kidney injury marker-NGAL, as up-regulated in both genotypes and SCD with and without proteinuria. Metascape-based analysis of biological processes revealed “Cellular component disassembly” associated with proteins expressed in SCD but not regulated between PU and no PU and “leukocyte chemotaxis” down-regulated in SCD_PU vs. SCD. Interestingly, “Integrin-cell-surface interactions” was associated with proteins up-regulated between SCD_PU vs. SCD which is consistent with endothelial hyperplasia in the setting of glomerular hyperfiltration. To investigate the effect secreted urine proteins have on human podocytes in vitro, immortalized podocytes supplemented with SCD_PU urine showed elevated p53 levels in both immunofluorescence staining and RT-PCR compared to SCD. Additionally, RT-PCR revealed elevated levels of VEGF, NGAL and the pro-inflammatory proteins-TGFβ, IL6, IL8 and TNFα.

**Conclusion:** We hypothesize that the increased number of endothelial cells in hyperplasia and hyperfiltration leads to more Integrin-mediated links to podocyte foot processes at the glomerular basement membrane and to glomerular fibrosis. Severe inflammation and kidney injury in SCD_PU patients is induced by the TGFβ-p53 axis.

## Introduction

Sickle cell disease (SCD) is a group of inherited blood disorders affecting hemoglobin, the protein in red blood cells (RBCs) that carries oxygen. SCD is primarily caused by a mutation in the beta-globin gene (*HBB*), which is responsible for producing hemoglobin [1]. This mutation results in the production of an abnormal form of hemoglobin, called hemoglobin S (HbS), instead of the normal hemoglobin A (HbA) [2], [3]. HbS leads RBCs to adopt a rigid, sickle-like shape and this deformed shape prevents the cells from easily passing through capillaries, leading to blockages and vaso-occlusion [4], [1]. SCD impacts multiple body systems, including kidneys, brain, liver, heart, gallbladder, eyes, bones, and joints, characterized by acute episodes of illness and gradual organ damage [4]. Repeated vaso-occlusive episodes and inflammation results in cumulative organ damage, which becomes more evident as individuals get older. Recurrent pain, fatigue, anemia, and increased infection susceptibility are some of the common symptoms manifested in SCD patients, while acute chest syndrome, delayed growth and development, stroke and organ damage are also reported [5], [6], [4]. SCD is more common among individuals from the Mediterranean, Middle East, South Asian, Caribbean, with the highest prevalence amongst individuals of African ancestry. It is also one of the most widespread severe single-gene disorders globally [7], [4].

SCD includes several types, with sickle cell anemia (SS) and sickle hemoglobin-C disease (SC) being among the most commonly observed. Patients with sickle cell anemia (HbSS) have homozygous HbS, inheriting one mutated beta-globin gene from each parent. In contrast, individuals with HbSC inherit a different mutation in their beta-globin genes, resulting in the production of both hemoglobin C and hemoglobin S [http://scancainc.org/learn/types-of-sickle-cell-disease/, accessed March 18 2025], [8], [9] [10]. Patients with an SS genotype have more severe clinical symptoms, while those with an SC genotype manifest milder symptoms [8], [9]

Nephropathy associated with SCD can lead to a range of renal complications including proteinuria, hyposthenuria, hematuria, renal papillary necrosis, renal tubular disorders, acute and chronic kidney injury, sickle cell glomerulopathy, and renal medullary carcinoma [11]. These kidney impairments often progress to chronic kidney disease (CKD), frequently requiring renal replacement therapy [1], [12], [13]. Findings from epidemiological studies also indicate an association between SCD and the occurrence of proteinuria and microalbuminuria in patients with autosomal dominant polycystic kidney disease (ADPKD) [12], [14], [15].

The pathophysiology of sickle cell nephropathy includes repeated sickling cycles favoured by hypoxic conditions in the renal medulla with vaso-occlusion in the vasa recta [11], [16], [17]. Ataga et al. revealed that prostaglandin release under hypoxic conditions leads to vasodilation and hyperfiltration [17] referring to reports of decreased hyperfiltration in SCD but not healthy control after administration of the prostaglandin inhibitor indomethacin [18]. Hyperfiltration among other factors such as glomerular hypertension and decreased nitric oxide [17] contributes to developing proteinuria although the detailed mechanisms are not fully elucidated.

Approximately 9.1% of the global population is affected by CKD [19]. CKD is linked to a wide range of adverse clinical outcomes, including cardiovascular disease, increased hospitalizations and death [20], [21], [19]. Regardless of the initial diagnosis, CKD shares common pathological mechanisms, including proteinuria, glomerular hyperfiltration, progressive renal scarring, and loss of kidney function [22]. Proteinuria is a condition where excess protein, primarily albumin, leaks into the urine due to kidney damage or dysfunction, disrupting the kidneys’ normal filtering process [23], [24]. Proteinuria can mediate inflammation and fibrosis, ultimately leading to kidney failure [24]. It is a well-known marker of kidney damage and is commonly associated with conditions such as CKD, diabetes, hypertension, and distinct immune disorders. Persistent proteinuria is a key indicator of severe CKD and can contribute to a decline in kidney function and increase the risk of cardiovascular disease [25]. Numerous studies have also revealed a correlation between proteinuria levels and an increased risk of cardiovascular disease [22]. Proteinuria is observed in approximately 8% to 33% of the general population [26], [27], [25]. Given that proteinuria and urinalysis abnormalities, and renal impairment were observed in African SCD patients, urinalysis abnormalities, kidney function assessment should be an active pursuit in patients with SCD [28]. Therefore, in this study we investigated novel urine-based kidney injury biomarkers as additional diagnostic tools for first line screening of SCD patients.

The aim of this study was to further elucidate the etiology of proteinuria in SCD and to identify SCD-associated kidney injury and inflammatory biomarkers based on analysing urinary proteins. We employed a Human XL cytokine and a kidney injury-specific biomarker array-based assay to analyze the expression and levels of urine-shed proteins from a cohort of SCD patients from Ghana. Some patients in the cohort were diagnosed with proteinuria during urine analysis. To validate our findings from the initial array analyses, we performed an ELISA-based quantitative protein analysis for Clusterin and NGAL. We compared SCD patients with and without proteinuria to healthy controls. The study involved urine analysis based on two approaches: pooled urine samples from healthy individuals, SCD patients without, and SCD patients with proteinuria, as well as a genotype-specific (SS, SC) analysis.

## Methods and materials

### Ethics Statement

Ethics approval for this current analysis was obtained locally from the Korle-Bu Teaching Hospital Institutional Review Board with approval protocol identification number KBTH-IRB/000261/2024. Written informed consent was obtained from all participants. Participants unable or unwilling to give consent or who were institutionalized were excluded.

### Participants

The current analysis included data from an initial project involving sickle cell disease patients. The initial study involved known SCD patients attending clinic at the Ghana Institute of Clinical Genetics (GICG) (Sickle Cell Clinic for adolescents and adults) in the Greater Accra Region of Ghana between February 2021 to May 2022. The institute attends to SCD patients 13 years and above and its environs on out-patient basis; approximately 50 patients are seen per day. It also receives referrals from other parts of the country. Consented persons with Sickle cell disease aged ≥18 years with and without clinical evidence of proteinuria or albumin/creatinine ratio ≥ 3.0 mg/mmol (30 mg/g) and healthy controls with no evidence of kidney disease were consecutively recruited.

The CKD-EPI equation without the race adjustment was used in adults and the Schwartz formula [29] in children aged less than 16 years. The following people were excluded from the study; pregnant females, smokers, history of recreational drug or alcohol abuse, current over-the-counter drug use, NSAID use within the last three months, persons with acute illness at time of enrollment, history of chronic illness, current evidence or history of renal disease and people who had sickle Cell crises within the last three months. Healthy persons without SCD or CKD were defined as individuals with eGFR 60 mL/min/1.73 m2 and albumin/creatinine ratio < 3.0 mg/mmol (<30 mg/g) and HB electrophoresis negative for sickle cell disease. Random urine samples were collected from cases and controls and aliquots of 10 mLs were taken into cryovials. All participants of this study are from African ancestry.

### Urine sample preparation and participant details

This analysis utilized urine samples collected from 10 healthy individuals and 20 patients from a cohort of individuals with sickle cell disease (SCD) in Ghana. Among the SCD patients, 10 were diagnosed with proteinuria and classified as the analysis group SCD with proteinuria (SCD_PU). The participants ranged in age from 18 to 49 years. Urine analysis was conducted using two approaches: pooled urine samples representing healthy individuals, SCD patients, and SCD_PU, as well as a genotype-specific (SS, SC) analysis. For each condition, urine samples from 10 individuals were pooled in equal volumes, and a final volume of 500 µl per condition was used for each of the array experiments. Clinical features of patients whose urine samples were pooled in the SCD_PU, SCD and control groups are listed in Table 1.

**Table 1:** Clinical features of patients whose urine samples were pooled into the SCD_PU, SCD and control groups.

| AGE | GENDER | Genotype | Std dipstick | Hb | Wbc | Platelet | S-Cr | GFR(CKC) | S- Alb. | U-Albu. | U-Cr | U-Al/Cr |
| --- | --- | --- | --- | --- | --- | --- | --- | --- | --- | --- | --- | --- |
| **SCD_PU** |  |  |  |  |  |  |  |  |  |  |  |  |
| 27 | M | SS | Positive | 6,8 | 14,9 | 518 | 31 | >89 | 36 | 809,61 | 5,88 | 137,69 |
| 21 | M | SC | Positive | 10,4 | 13,3 | 258 | 72 | >89 | 36 | 33,25 | 13,69 | 2,43 |
| 21 | F | SS | Positive | 9,9 | 8,2 | 538 | 64 | >89 | 40 | 947,21 | 9,58 | 98,87 |
| 29 | F | SS | Positive | 9,5 | 7,9 | 490 | 28 | >89 | 36 | 176,33 | 6,29 | 28,03 |
| 28 | F | SS | Positive | 6,9 | 9,9 | 461 | 38 | >89 | 47 | 314 | 0,49 | 640,82 |
| 31 | F | SC | Positive | 9,9 | 11,5 | 460 | 65 | >89 | 43 | 333,15 | 8,69 | 38,34 |
| 19? | M | SC | Positive | 9,5 | 16,3 | 655 | 61 | >89 | 36 | 246,74 | 18,14 | 13,6 |
| 26 | F | SS | Positive | 6,7 | 6,2 | 358 |  |  |  |  |  |  |
| 28? | F | SS | Positive | 6,5 | 12,6 | 616 | 71 | >89 | 42 | 885,78 | 16,63 | 53,26 |
| **SCD** |  |  |  |  |  |  |  |  |  |  |  |  |
| 18 | M | SC | Negative | 8,7 | 8,4 | 480 | 64 | >89 | 44 | 36,97 | 7,09 | 5,21 |
| 45 | F | SS | Negative | 8 | 9,1 | 257 | 46 | >89 | 44 | 20,15 | 8,49 | 2,37 |
| 48/49 | F | SS | Negative | 9,5 | 6,1 | 88 | 60 | >89 | 48 | 5,86 | 9,8 | 0,6 |
| 18 | M | SC | Negative | 10,4 | 14,4 | 412 | 55 | >89 | 43 | 17,52 | 4,16 | 4,21 |
| 42 | F | SC | Negative | 11,8 | 7,5 | 272 | 103 | 72 | 40 | 3,5 | 6,91 | 0,51 |
| 37 | F | SS | Negative | 11,2 | 7,07 | 242 | 55 | >89 | 45 | 3,88 | 8,32 | 0,47 |
| 43 | F | SS | Negative | 10,2 | 5,64 | 305 | 30 | >89 | 46 | 5,56 | 5,39 | 1,03 |
| 23? | F | SS | Negative | 6,8 | 8,2 | 386 | 37 | >89 | 46 | 53,98 | 19,56 | 2,76 |
| 29 | F | SS | Negative | 8 | 7,2 | 436 | 53 | >89 | 44 | 33,81 | 11,05 | 3,06 |
| **control** |  |  |  |  |  |  |  |  |  |  |  |  |
| 32 | M | AA | Negative | 14,2 | 4,3 | 276 | 120 | 69 | 42 | 0,97 | 10,62 | 0,09 |
| 21 | M | AC | Negative | 14,8 | 7,5 | 223 | 55 | >89 | 46 | 2,84 | 9,94 | 0,29 |
| 31 | F | AC | Negative | 13,4 | 5,7 | 272 | 60 | >89 | 41 | 5,23 | 19,75 | 0,26 |
| 34 | F | AA | Negative | 13 | 6,2 | 86 | 58 | >89 | 44 | 4,24 | 12,54 | 0,34 |
| 31 | M | AA | Negative | 15,2 | 3,9 | 251 | 78 | >89 | 47 | 4,29 | 16,48 | 0,26 |
| 39 | M | AA | Negative | 14,6 | 5,1 | 189 | 75 | >89 | 41 | 4,6 | 10,33 | 0,45 |
| 43 | M | AA | Negative | 13,8 | 4,7 | 237 | 71 | >89 | 42 | 68,52 | 14,56 | 4,71 |
| 38 | M | AA | Negative | 14,1 | 3,9 | 149 | 71 | >89 | 45 | 13,15 | 15,37 | 0,86 |
| 30 | M | AA | Negative | 14,3 | 6,2 | 259 | 82 | >89 | 40 | 43,5 | 14,94 | 2,91 |
| 27 | F | AA | Negative | 12,8 | 9,2 | 209 | 53 | >89 | 43 | 4,88 | 12,21 | 0,4 |

### Array-based cytokine and kidney injury biomarker analysis

As with our previous studies using urine samples from AKI and CKD patients [30], [31], the current study used pooled urine samples from healthy individuals, SCD and SCD.PU patients, as well as pools of genotype-specific samples (SS, SC). These were hybridized onto the Human Kidney Biomarker Array (Human Kidney Biomarker Array Kit, Catalog Number ARY019) and the Proteome Profiler Array (Human XL Cytokine Array Kit, Catalog Number ARY022B) from Research And Diagnostic Systems, Inc. (Minneapolis, MN, USA), following the manufacturer’s protocol. Images of the hybridized arrays were captured using the Fusion FX instrument (PeqLab, Erlangen, Germany). The detailed workflow of the analyses is depicted in Supplementary Figure 1.

### Image analysis of cytokine arrays

The images of the hybridized cytokine assays were imported into the FIJI/ImageJ software [32] for preprocessing, grid-finding and quantification via the FIJI Microarray Profile plugin by Bob Dougherty and Wayne Rasband (https://www.optinav.info/MicroArray_Profile.htm, accessed on 06 January 2025). Details of the image analysis including semi-automatically grid-finding employing an R script for interpolation of missing spots are described in our previous publication [31].

### Array data analysis

Quantification results from the microarray plugin within FIJI/ImageJ were imported into the R/Bioconductor [33] environment where they were normalized with the Robust Spline Normalization from the R/Bioconductor package lumi [34]. Up-and down-regulated cytokines for the comparisons SCD vs. control, SCD_PU vs. control, SCD_PU vs SCD and SS vs SC were determined using the criteria: detection-p-value in at least one case < 0.05, p-value from the limma [35] R package < 0.05, false-discovery-rate < 0.25, ratio >1.5 (up-regulation) or ratio < 0.6667 (down-regulation). Hierarchical cluster analysis dendrograms and heatmaps were drawn via the heatmap.2 function from the R package gplots [36] using complete linkage as cluster agglomeration method and the Pearson correlation coefficient as similarity measure. Barplots with error bars indicating standard error of the mean were generated via the R methods barplot and arrows.

### Enzyme-linked Immunosorbent assay (ELISA)

Secreted Clusterin and Lipocalin-2 (NGAL) detected in pooled urine samples of SCD and SCD_PU patients using the DuoSet ELISA (R&D Systems, Minneapolis, MN, USA) was performed as described by the manufacturer. Optical density was measured using the EPOCH2 spectrophotometer (BioTek, Winooski, Vermont, USA) at 450 nm with wavelength correction at 540 nm. 4-PL curve fitting was performed to calculate concentrations.

### Cell culture

The human SV40, temperature sensitive podocyte immortal cell line (AB 8/13) [37] proliferates at 33°C, 5% CO2 on plastic when cultured in RPMI 1640 (Gibco®, life technologies, California) supplemented with 10% fetal bovine serum (Gibco®,) and 1% Penicillin and Streptomycin (Gibco®,).

For differentiation into podocytes we followed our previously published protocols [38], [39]. In brief, the cells were seeded on Corning® Collagen type I (Merck, Darmstadt, Deutschland) coated plates and cultured in the differentiation medium-Advanced RPMI 1640 (Gibco®) supplemented with 10% fetal bovine serum, 1% Penicillin and Streptomycin, and 30 μM retinoic acid (Sigma-Aldrich Chemistry, Steinheim, Germany), incubated at 37 °C, 5% CO2 for 14 days. The medium was changed once a week. After 14 days the podocytes were treated with for 5 days with 30% urine from healthy individuals, sickle cell patients without proteinuria (SCD) and sickle cell patients with proteinuria SCD_PU for 5 days.

### Immunofluoresence-based protein staining

Human immortalized podocytes (AB 8/13) were fixed with 4% paraformaldehyde for 15 min at room temperature (RT). Cells were washed three times with PBS. Afterwards an incubation with 0.05% Tween-20/PBS for 10 minutes. The cells were treated three times with PBS and continued blocking the cells with 3% BSA/PBS. Thereafter they were incubated with the respective primary antibody (Cell signaling; 2527S) diluted 1:200 overnight at 4 °C. Followed by one washing step with Triton x-100/PBS and two washing steps with PBS. The cells were incubated for 1 h with Alexa488-conjugated secondary antibody diluted 1:500 and nuclear Hoechst 1:5000(Thermo Fisher; H3569) at RT. Fluorescence images were captured by a LSM700 microscope (Carl Zeiss).

### RNA isolation

For RNA isolation from the human immortalized podocyte cell (AB 8/13) the ZYMO Research Kit Direct-zol™ RNA Miniprep R20 was utilized following the manufacturer’s protocol. Cells were detached by adding Tryple E and incubated for 5 min. Afterwards, a centrifuging step of 4 mins 1000 g and the cell pellets processed following manufactureŕs protocol. The RNA was eluted by adding 35 μL of DNase/RNase-free water to the column and centrifuging for 1 minute at 16,000 g. The eluted RNA samples were immediately placed on ice, and concentrations measured with a Nanodrop device.

### Real-Time PCR based gene expression quantification of pro-inflammatory and kidney injury-associated genes

RNA samples were run in triplicate on a 384-well reaction plate with Power Sybr Green PCR Master Mix (Applied Biosystems, Foster City, CA, USA) using Step One Plus Real-Time PCR systems. The amplification conditions were denaturation at 95 ^0^C for 13 min followed by 35 cycles of 95 ^0^C for 50 s, 60 ^0^C for 45 s, and 72 ^0^C for 30 s. To normalize the quantitative real-time PCR, the ribosomal encoding gene-RPL0 was used. The primer sequences are listed in Supplementary Table S1. The results were analyzed using the 2−ΔΔCT method and specified by fold change expression. The samples were run in independent experiments three times. The mean value of these experiments was used for the calculations. The statistical significance was calculated using the Two-Sample test assuming unequal variances [40] with a significance threshold p = 0.05.

## Results

### Comparison of kidney injury markers between SCD with and without Proteinuria

We pooled urine samples from SCD_PU, SCD without proteinuria and healthy controls and analyzed these on the Human Kidney Biomarker Array. Figure 1A shows the cluster dendrogram with separate clusters of SCD with PU (SCD_PU), SCD without PU (SCD) and healthy control (Ctrl). The analysis revealed distinct kidney protein signatures of SCD_PU, SCD and ctrl. The heatmaps in Figure 1B-D distinguish expression of differentially expressed proteins between SCD and Ctrl (B), between SCD_PU and Ctrl (C) and between SCD_PU and SCD (D). Resistin, IL1RA, RAGE, TFF3 and uPA (PLAU) are up-regulated in SCD_PU vs SCD while DPPIV, FABP1, Angiotensinogen, PSA (KLK3), MMP9, Adiponectin, ANPEP and Clusterin are down-regulated. Clusterin had lower expression in SCD_PU than in SCD, in line with reports of Clusterin attenuating kidney disease [41]. We therefore followed-up this protein with ELISA measurements which confirmed higher expression of Clusterin in SCD than in control but lower expression in SCD_PU than in control (Figure 1E). Figure 1F provides a global view of the expression of all proteins under the three conditions on the Kidney Injury Array.

**Figure 1:**
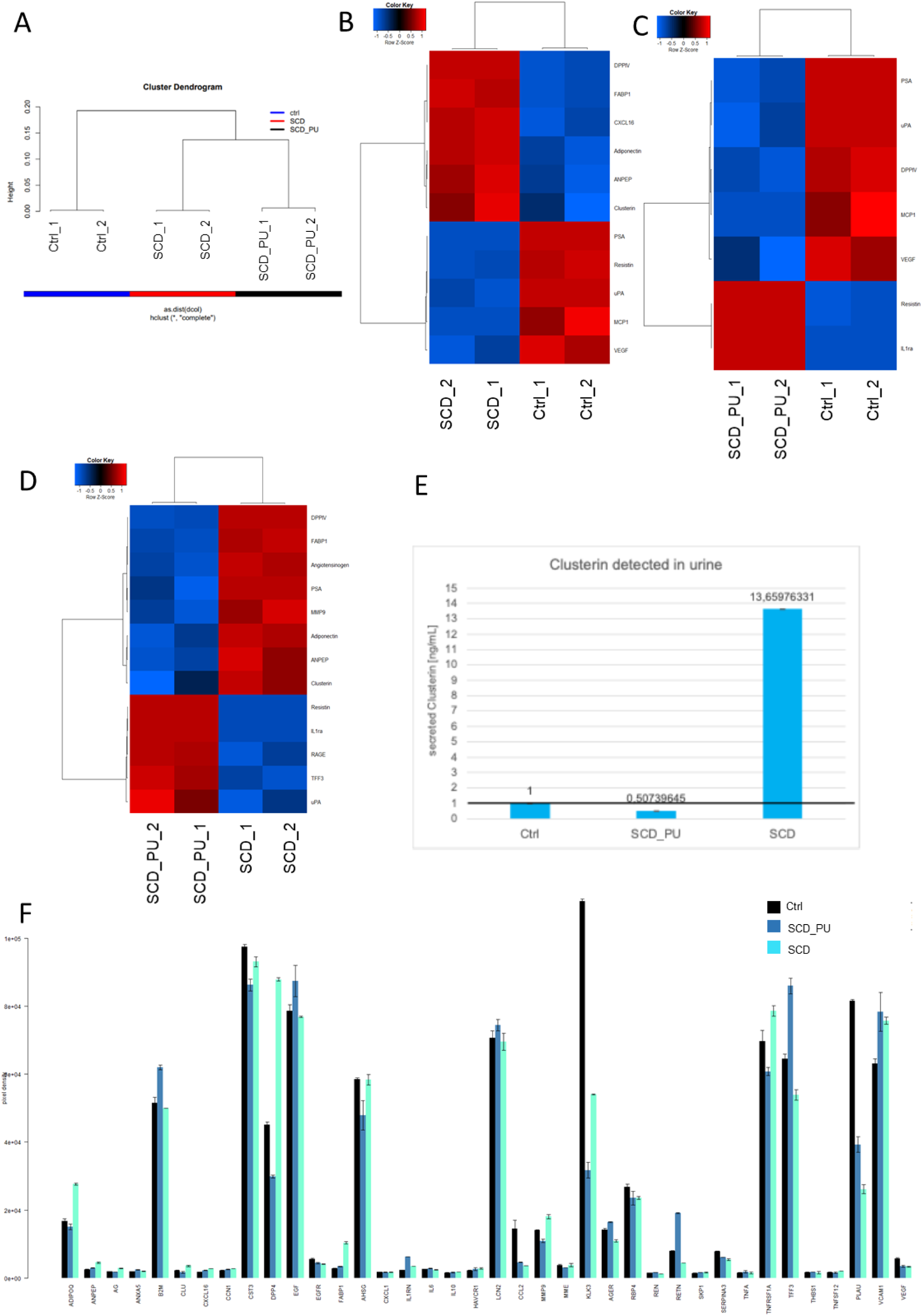
Pooled urine samples of SCD with and without Proteinuria (PU) have distinct kidney protein signatures. Pooled urine samples of SCD with and without Proteinuria were analyzed on the Human Kidney Biomarker Array. (A) The cluster dendrogram shows separate clusters of SCD with PU (SCD_PU), SCD without PU (SCD) and healthy control (Ctrl). (B) The heatmap shows expression of differentially expressed proteins between SCD and Ctrl. (C) The heatmap shows expression of differentially expressed proteins between SCD_PU and Ctrl. (D) The heatmap shows expression of differentially expressed proteins between SCD_PU and SCD. Resistin, IL1RA, RAGE, TFF3 and uPA (PLAU) are up-regulated in SCD_PU while DPPIV, FABP1, Angiotensinogen, PSA (KLK3), MMP9, Adiponectin, ANPEP and Clusterin are down-regulated. (E) The bar chart of ELISA measurements confirms higher expression of Clusterin in SCD than in control but lower expression in SCD_PU than in control. (F) The bar plot shows the absolute expression of all proteins on the Human Kidney Biomarker Array.

### Comparison of kidney injury markers between SCD with and without Proteinuria

Furthermore, we set out to compare expression of cytokines between SCD_PU, SCD and healthy control. Again, we pooled urine samples of SCD_PU, SCD and control and analyzed them - this time on a Human Cytokine XL array. The heatmaps in Figure 2A and B show expression of differentially expressed proteins between SCD and ctrl and between SCD_PU and ctrl. The heatmap in Figure 2C shows expression of differentially expressed proteins between SCD_PU and SCD. Emmprin, IL1RA, SerpinE1, SHBG, Osteopontin, CD31, IL-18-Bpa and IL-17A are up-regulated in SCD_PU while RBP4, DPPIV, Vitamin-D-BP, Complement-Component-C5-C5a, BAFF, MCP-1, IL-8 and M-CSF are down-regulated. The bar plot in Figure 2D provides an overview of the absolute expression of all proteins on the Human XL cytokine Array for the three conditions Ctrl, SCD_PU and SCD.

**Figure 2:**
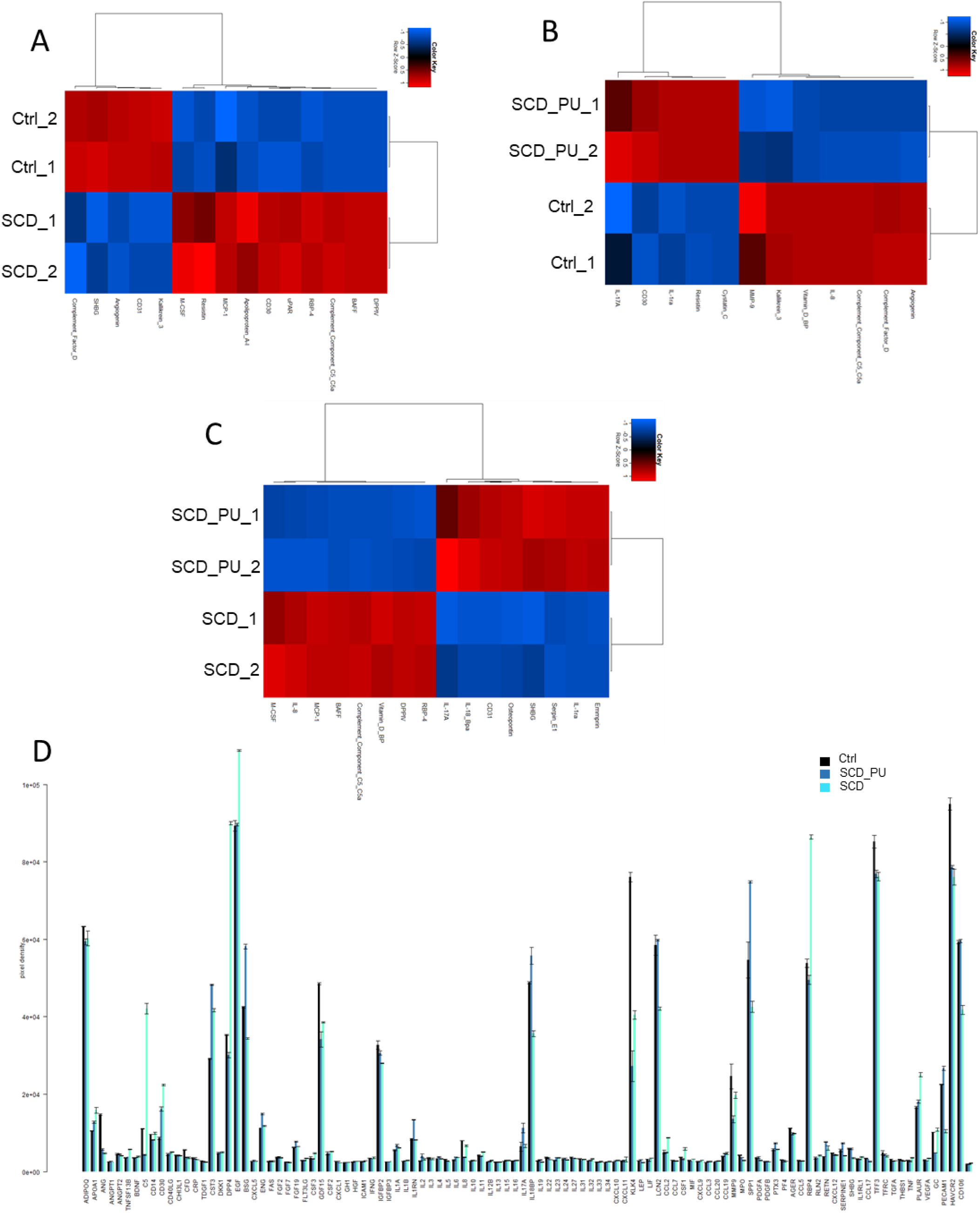
Pooled urine samples of SCD with and without Proteinuria (PU) have distinct cytokine signatures. Pooled urine samples of SCD with and without Proteinuria were analyzed on a Human XL cytokine array. (A) The heatmap shows expression of differentially expressed proteins between SCD and ctrl. (B) The heatmap shows expression of differentially expressed proteins between SCD_PU and Ctrl. (C) The heatmap shows expression of differentially expressed proteins between SCD_PU and SCD. Emmprin, IL1RA, SerpinE1, SHBG, Osteopontin, CD31, IL-18-Bpa and IL-17A are up-regulated in SCD_PU while RBP4, DPPIV, Vitamin-D-BP, Complement-Component-C5-C5a, BAFF, MCP-1, IL-8 and M-CSF are down-regulated. (D) The bar plot shows the absolute expression of all proteins on the Human XL cytokine Array.

In order to assign functional annotations associated with the development of proteinuria in SCD we performed a Metascape analysis of up-, down-and not-regulated cytokines by comparing SCD_PU vs SCD (Figure 3). Figure 3A depicts the Reactome pathways “Extracellular matrix organization” and “Integrin cell surface interactions” over-represented in cytokines up-regulated in SCD_PU. Figure 3B lists the Gene Ontologies “negative regulation of leukocyte chemotaxis”, “regulation of leukocyte chemotaxis” and “regulation of granulocyte chemotaxis” over-represented in cytokines down-regulated in SCD_PU. The gene ontology “cellular component disassembly” was over-represented in not-regulated cytokines but expressed in both conditions (Figure 3C). Figure 3D shows a heatmap and Figure 3E a protein interaction network resulting from the Metascape analysis of up-, down-and not-regulated cytokines.

**Figure 3:**
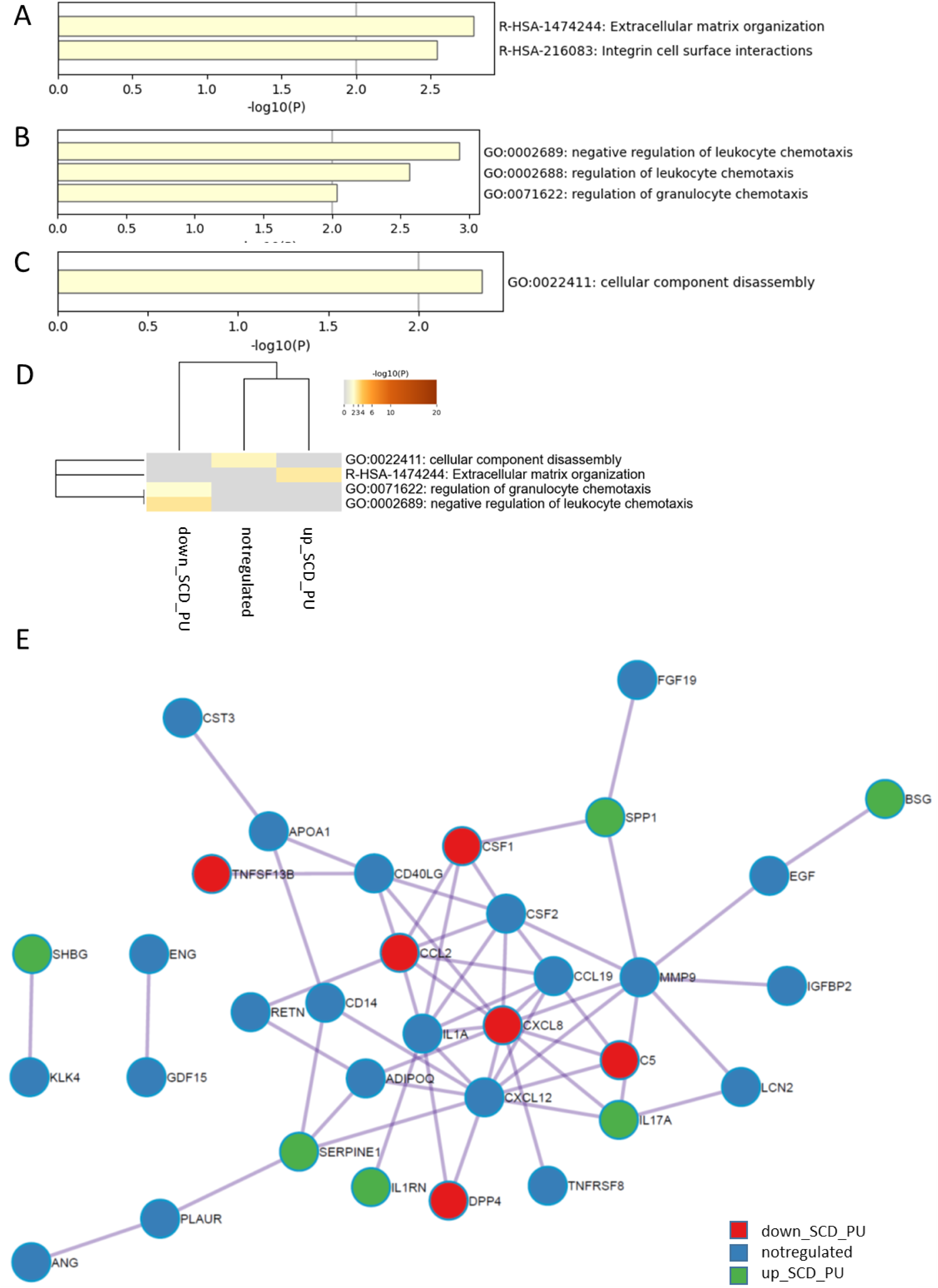
**SCD_PU vs SCD Metascape analysis**. Up-, down-and not-regulated cytokines when comparing SCD_PU vs. SCD were subjected to metascape analysis. (A) Reactome pathways “Extracellular matrix organization” and “Integrin cell surface interactions” were over-represented in cytokines up-regulated in SCD_PU. (B) Gene ontologies “negative regulation of leukocyte chemotaxis”, “regulation of leukocyte chemotaxis” and “regulation of granulocyte chemotaxis” were over-represented in cytokines down-regulated in SCD_PU. (C) The gene ontology “cellular component disassembly” was over-represented in cytokines not-regulated but expressed in both conditions.(D) shows a heatmap and (E) a protein interaction network resulting from the combined Metascape analysis of up-, down-and not-regulated cytokines.

### Comparison of Proteinuria in genotypes SS and SC

Pooled urine samples of genotypes SS and SC from SCD patients with and without Proteinuria were analyzed on Human XL cytokine and kidney biomarker arrays (Figure 4). The heatmaps in Figure 4A and B show expression of differentially expressed kidney injury marker proteins (Figure 4A) and cytokines (Figure 4B) between SCD_PU patients with the genotypes SS and SC. The bar plots in Figure 4C and D show expression of differentially expressed kidney injury marker proteins (Figure 4C) and cytokines (Figure 4D) between SCD_PU patients with the genotypes SS and SC. The kidney injury markers Lipocalin-2 (LCN2 / NGAL), MMP9, CCN1 and IL1RN (IL1RA) are up-regulated in SS_PU while ANPEP, CLU (Clusterin), IL6, KLK3, RETN and SERPINA3 are down-regulated in SS_PU (Figure 4C). Amongst the up-regulated cytokines in SS_PU (Figure 4D), LCN2 and MMP9 levels were also upregulated in the kidney injury analyses. (E) Up-regulation of the established kidney injury marker LCN2 (NGAL) between SS_PU and SC_PU (C) was confirmed by ELISA measurements in SCD_PU vs SCD (Figure 4E) and between SS_PU and SC_PU (Figure 4F).

**Figure 4:**
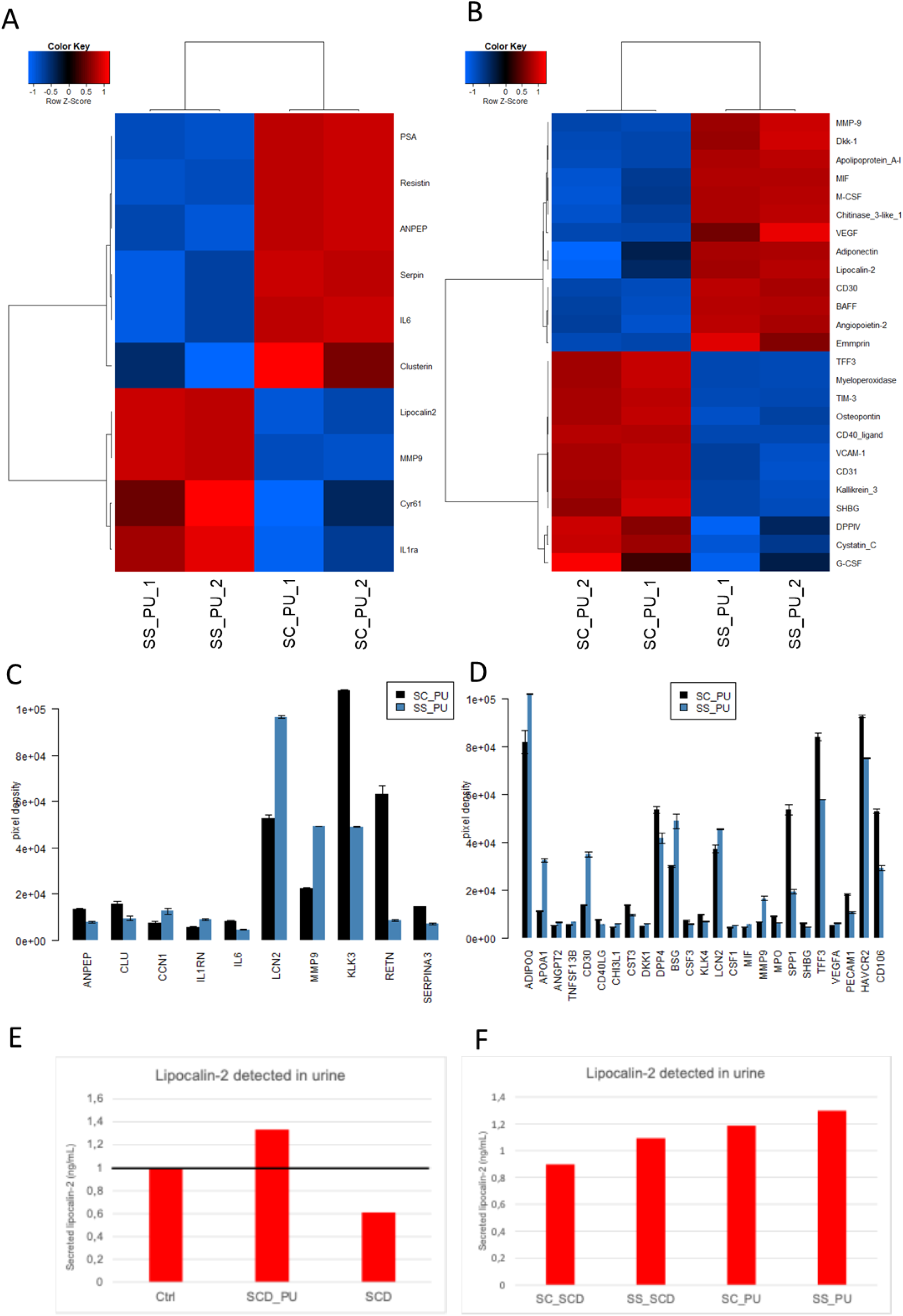
Comparison of Proteinuria in genotypes SS and SC on Human cytokine XL and kidney arrays. Pooled urine samples of genotypes SS and SC from SCD patients with and without Proteinuria were analyzed on the Human XL cytokine and kidney biomarker arrays. (A) The heatmap shows expression of differentially expressed kidney injury marker proteins between SCD_PU patients with the genotypes SS and SC. (B The heatmap shows expression of differentially expressed cytokines between SCD_PU patients with the genotypes SS and SC. (C) The bar plot shows expression of differentially expressed kidney injury marker proteins between SCD_PU patients with the genotypes SS and SC. Lipocalin-2 (LCN2 / NGAL), MMP9, CCN1 and IL1RN (IL1RA) are up-regulated in SS_PU while ANPEP, CLU (Clusterin), IL6, KLK3, RETN and SERPINA3 are down-regulated in SS_PU. (D) The bar plot shows expression of differentially expressed cytokines between SCD_PU patients with the genotypes SS and SC. Amongst the cytokines up-regulated in SS_PU, the increased levels of LCN2 and MMP9 were also confirmed in the kidney biomarker array. (E) Up-regulation of the established kidney injury marker LCN2 (NGAL) between SS_PU and SC_PU (C) was confirmed by ELISA in SCD_PU vs SCD and between SS_PU and SC_PU (F).

### The effects of urine-secreted cytokines from SCD, SCD_PU and healthy individuals on human podocytes

For this investigation, human immortalized podocytes (AB8/13) were cultured for 14 days in advanced RPMI medium supplemented with 30 µM retinoic acid (RA) on collagen type I-coated plates. The cells were then exposed for an additional five days with pooled urine samples from healthy controls, SCD and SCD_PU patients Following the treatment, light microscopy revealed no morphological changes in podocytes exposed to urine from all groups (Figure 5A). The podocyte specific marker Podocin (NPHS2) served as a positive control demonstrating successful podocyte differentiation whilst Phalloidin served to show the morphology of the cytoskeleton (Figure 5B). Immunofluorescence staining showed increased TP53 protein expression in podocytes treated with SCD_PU urine, thus implying activation of cell cycle arrest due to stress responses (Figure 5B). Quantitative real-time PCR analysis demonstrated a downregulation in the expression of the podocyte-specific markers Nephrin (NPHS1) and upregulation of CD2-associated protein (CD2AP). Both mRNA and protein levels of TP53 were upregulated (Figure 5C). Furthermore, transcripts of kidney injury markers-Cystatin C (CysC) and Neutrophil gelatinase-associated lipocalin (NGAL), and pro-inflammatory markers-Tumor necrosis factor alpha (TNF-α), Transforming growth factor beta (TGF-β), Vascular endothelial growth factor (VEGF), Interleukin 6 (IL-6) and Interleukin (IL-8) were significantly elevated in podocytes treated with SCD_PU urine, thus indicating enhanced cellular stress and injury. Clusterin (CLU) and Dipeptidyl peptidase-4 inhibitor (DPP4) levels were downregulated in both SCD and SCD_PU compared to the healthy control (Figure 5C).

**Figure 5:**
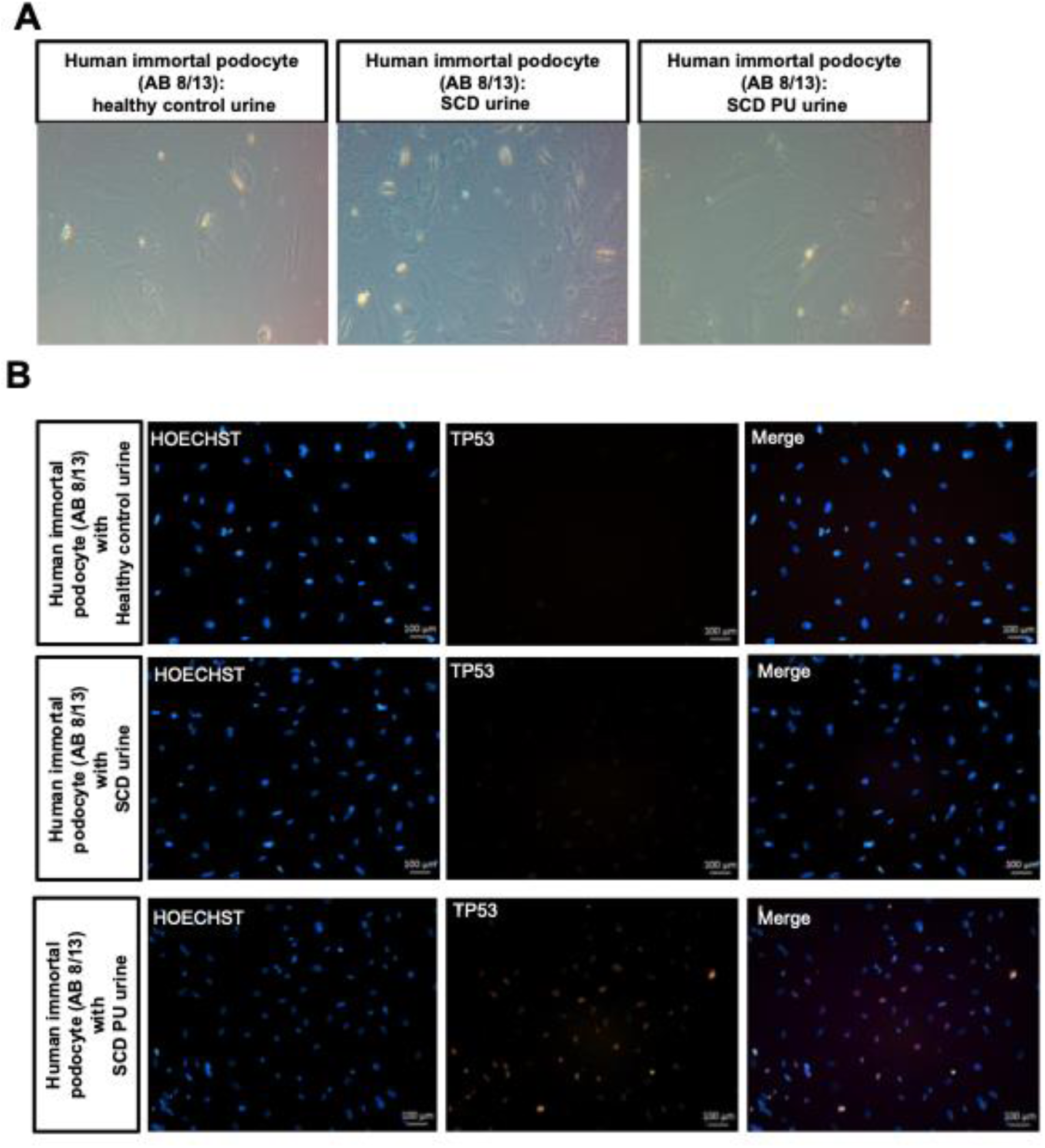
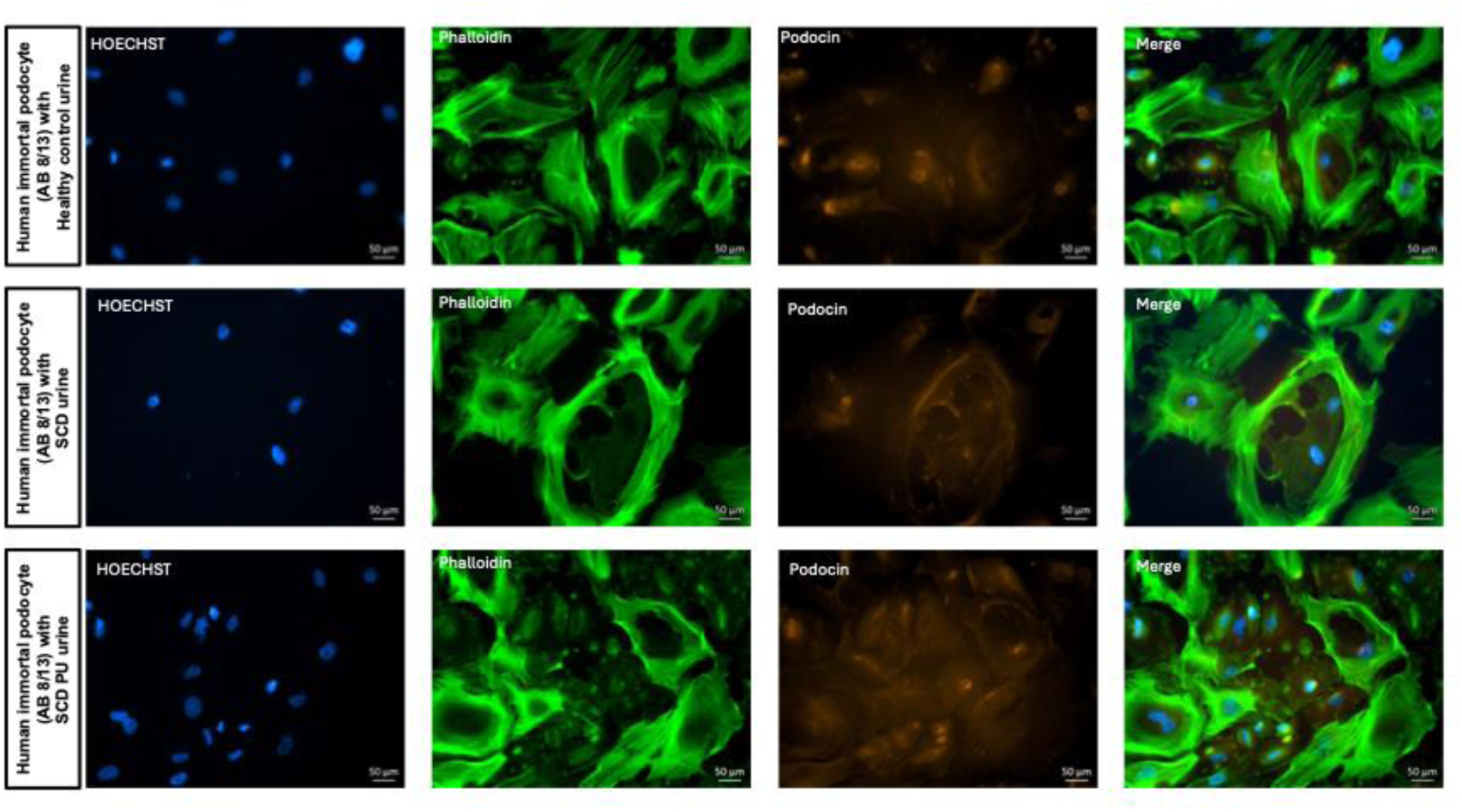
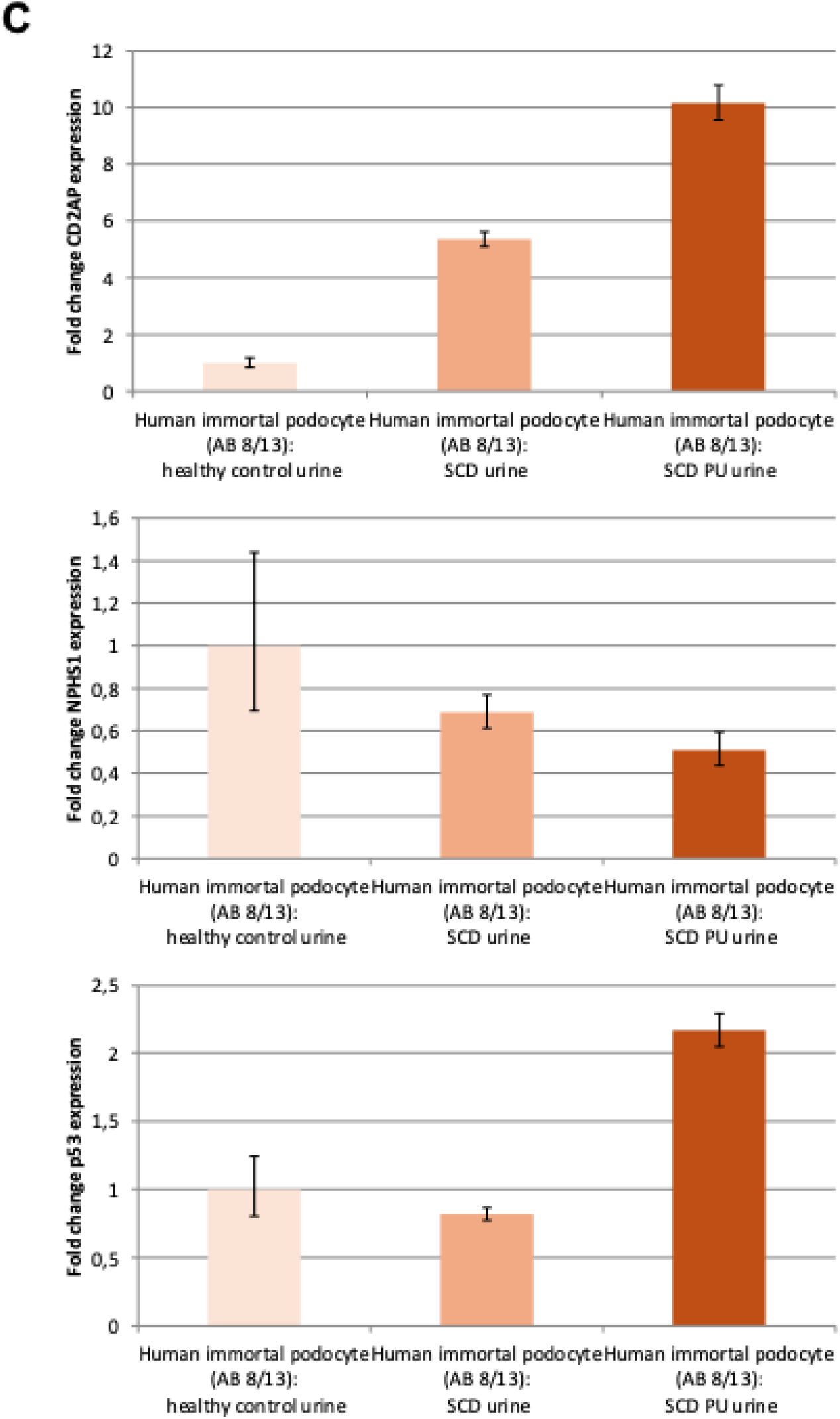
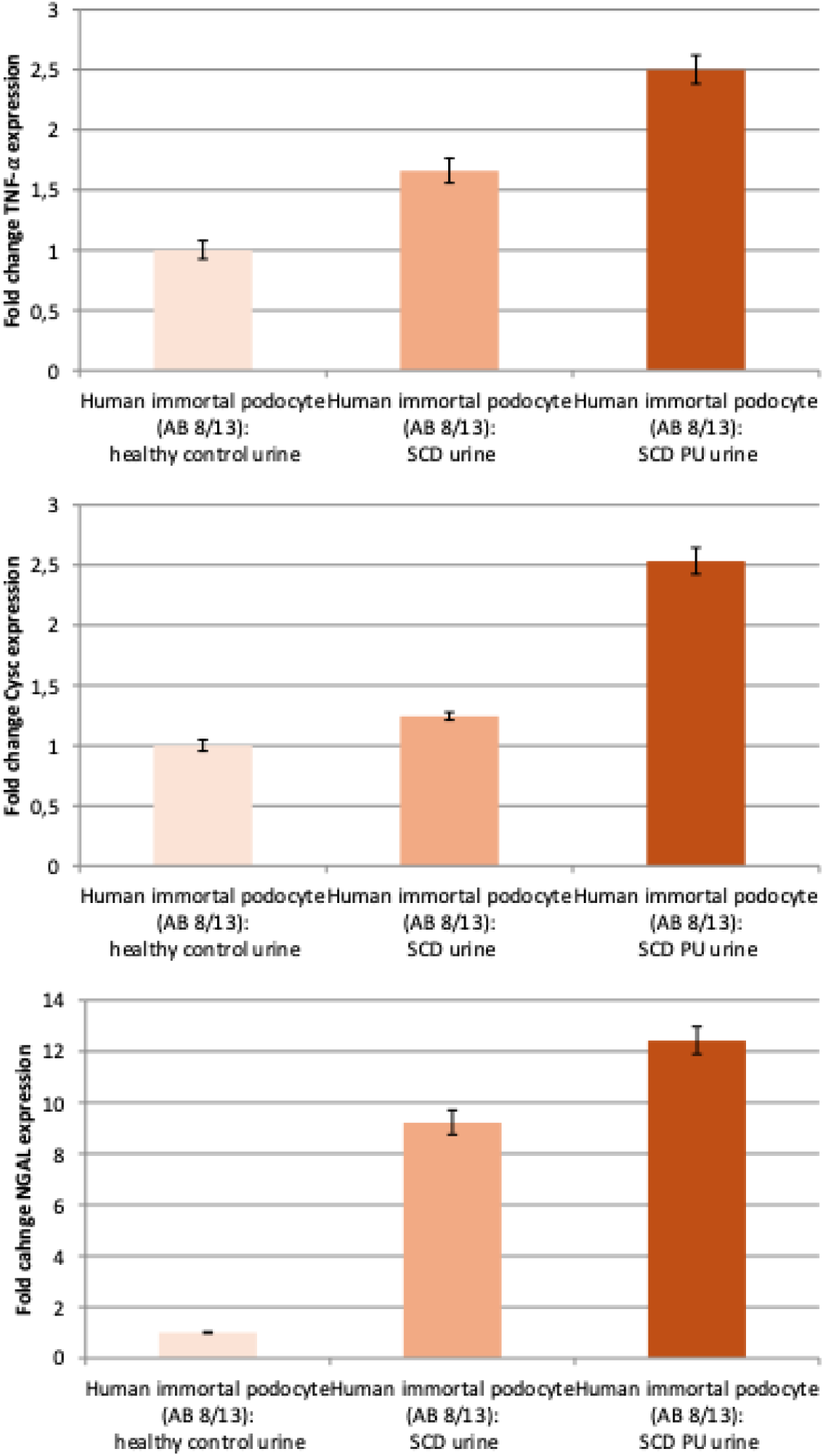
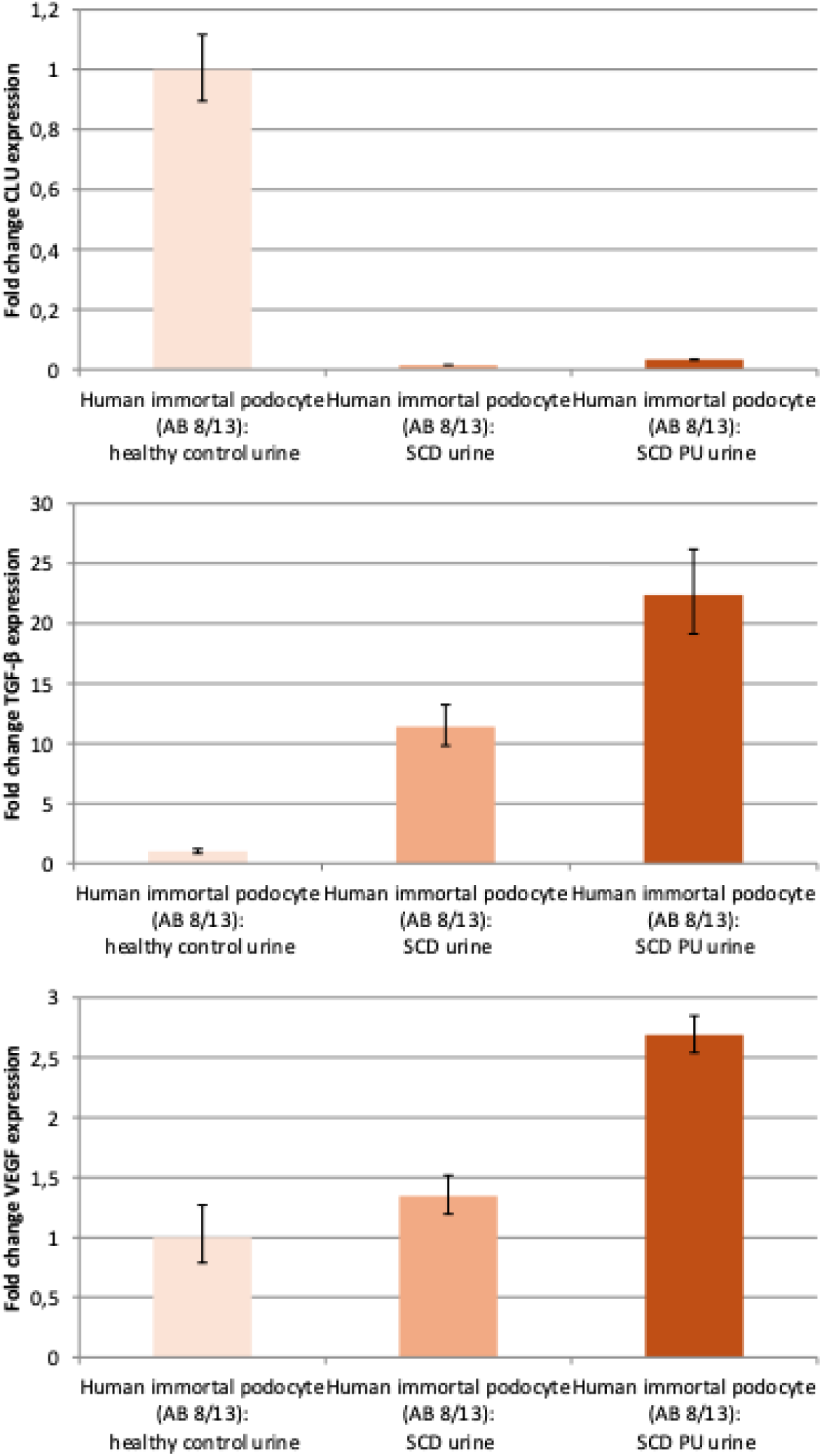
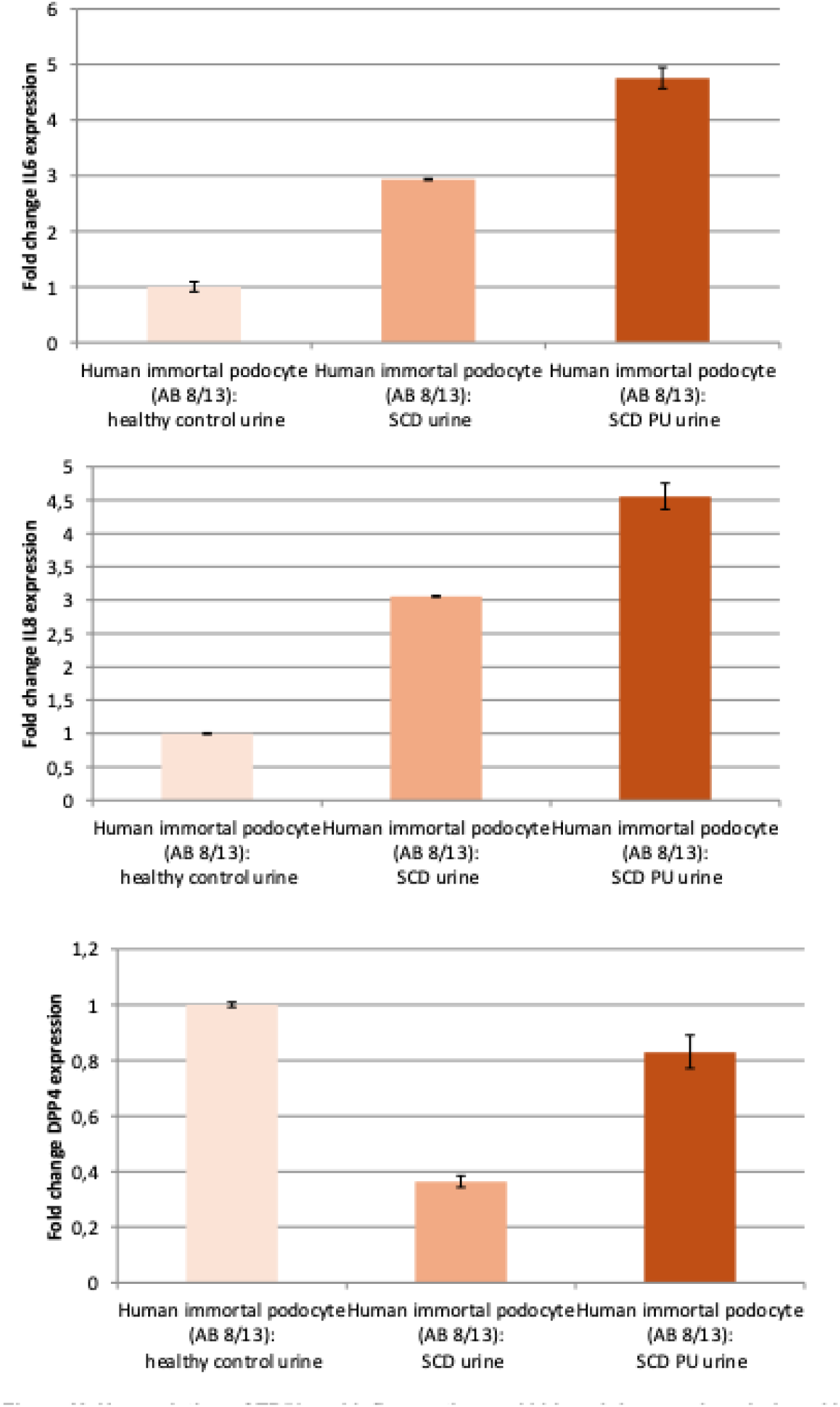
Upregulated expression of TP53, pro-inflammatory and kidney injury-associated markers in podocytes exposed to urine from SCD and SCD_PU patients. Human immortalized podocytes (AB8/13) were cultured at 37 °C for 14 days in Advanced RPMI medium supplemented with 30 µM retinoic acid (RA) on collagen type I-coated plates. The cells were then treated for five days with urine samples from healthy controls, SCD amd SCD_PU patients. (A) Representative light microscopy images of podocytes after five days of urine exposure. (B) TP53, NPHS2 and Phalloidin protein expression visualized by immunofluorescence staining. (C) Quantitative real-time PCR analysis of mRNA expression levels for podocyte markers (NPHS1, CD2AP), the regulator of cellular stress (TP53), and kidney injury-associated genes (CysC, NGAL and CLU) and pro-inflammatory associated genes (TNF-α, TGF-β, VEGF, IL6, IL8 and DPP4).

## Discussion

In previous studies we used urine samples to identify cytokines and kidney injury-associated biomarkers in patients diagnosed with Acute Kidney injury and Chronic kidney disease [31], [30].

In this study we investigated the similarities and differences between SCD_PU with proteinuria, SCD without proteinuria and healthy individuals with respect to the secretion of cytokines and kidney injury-associated proteins in pooled urine samples. Table 2 provides an overview of the resulting proteins.

**Table 2:** Up-and down-regulated proteins between SCD_PU, SCD and control and SS_PU and SC_PU.

| protein | SCD_PU_vs_SCD | SCD_PU_vs_ctrl | SCD_vs_ctrl | SS_PU_vs_SC_PU | array |
| --- | --- | --- | --- | --- | --- |
| ADIPOQ | down |  | up |  | kidney |
| ANPEP | down |  | up | down |  |
| AG | down |  |  |  |  |
| CLU | down |  | up | down |  |
| DPP4 | down | down | up |  |  |
| FABP1 | down |  | up |  |  |
| MMP9 | down |  |  |  |  |
| KLK3 | down |  | down | down |  |
| CCL2 |  | down | down |  |  |
| PLAU | up | down | down |  |  |
| VEGF |  | down | down |  |  |
| RETN | up | up | down | down |  |
| IL1RN | up | up |  |  |  |
| AGER | up |  |  |  |  |
| TFF3 | up |  |  |  |  |
| CXCL16 |  |  | up |  |  |
| IL6 |  |  |  | down |  |
| SERPINA3 |  |  |  | down |  |
| TNFSF13B | down |  | up | up | human_XL |
| C5 | down | down | up |  |  |
| DPP4 | down |  | up | down |  |
| IL8 | down | down |  |  |  |
| CCL2 | down |  | up |  |  |
| CSF1 | down |  | up | up |  |
| RBP4 | down |  | up |  |  |
| GC | down | down |  |  |  |
| ANG |  | down | down |  |  |
| CFD |  | down |  |  |  |
| KLK4 |  | down | down | down |  |
| MMP9 |  | down |  | up |  |
| CFD |  |  | down |  |  |
| SHBG | up |  | down | down |  |
| PECAM1 | up |  | down | down |  |
| BSG | up |  |  |  |  |
| IL1RN | up | up |  |  |  |
| IL17A | up | up |  |  |  |
| IL18BP | up |  |  |  |  |
| SPP1 | up |  |  | down |  |
| SERPINE1 | up |  |  |  |  |
| CD30 |  | up |  |  |  |
| CST3 |  | up |  | down |  |
| RETN |  | up | up |  |  |
| APOA1 |  |  | up | up |  |
| CD30 |  |  | up | up |  |
| CD40LG |  |  |  | down |  |
| CSF3 |  |  |  | down |  |
| MPO |  |  |  | down |  |
| TFF3 |  |  |  | down |  |
| HAVCR2 |  |  |  | down |  |
| CD106 |  |  |  | down |  |
| ADIPOQ |  |  |  | up |  |
| ANGPT2 |  |  |  | up |  |
| CHI3L1 |  |  |  | up |  |
| DKK1 |  |  |  | up |  |
| BSG |  |  |  | up |  |
| LCN2 |  |  |  | up |  |
| MIF |  |  |  | up |  |
| VEGFA |  |  |  | up |  |

Resistin, IL1RA, RAGE, TFF3 and uPA (PLAU) are up-regulated in SCD_PU vs SCD while DPPIV, FABP1, Angiotensinogen, PSA (KLK3), MMP9, Adiponectin, ANPEP and Clusterin are down-regulated.

Clusterin which we identified to be secreted at a lower level in SCD_PU than in SCD patients has been reported as biomarker for microalbinuria in a consecutive cohort study of 146 type 2 diabetes mellitus (T2DM) patients [42] and additionally attenuates renal fibrosis in rodent models [41]. Numerous studies have described an association between elevated Resistin and Proteinuria [43], [44], thus arguing that the level of Resistin is rather connected to inflammation and glomerular filtration impairment than to Insulin Resistance. The reported increased levels of Resistin corroborates our finding of up-regulated levels of Resistin in SCD_PU.

IL-1Ra is a receptor antagonist of the inflammation-mediator IL-1, and the balance between IL-1 and IL-1Ra plays an important role in several diseases including renal diseases [45]. IL-1 is involved in inflammation-associated development of diseases which can be reversed by the antagonist IL-1Ra– this has been exploited in therapies based on recombinant IL-1Ra (anakinra) [46]. Human immune cells can secrete IL-1Ra to compensate for IL-1-mediated inflammation. Thus, IL-1Ra can also be a marker of diseases. This has been shown in rat studies of glomerulonephritis with anti-glomerular basement membrane (GBM) antibodies showing production of IL-1 and IL-1Ra within the rat kidney [47]. Descamps-Latscha described elevated levels of IL-1Ra in patients at various stages of chronic renal failure and the correlation of IL-I Ra with the soluble TNFA-receptor,-TNF-sR55 [48].

The receptor of advanced glycation end product (RAGE) was up-regulated in SCD_PU. Increased levels of RAGE has been shown to induce oxidative stress in diabetic kidney disease and ageing kidney via NAPDH (nicotinamide adenine dinucleotide phosphate) oxidase [49], [50], [51]. In their review, Duni et al. described models of podocyte injury induced by increased levels of reactive oxygen species (ROS) which led to ultrastructural changes in podocytes, defective anchorage due to alteration in the adhesion molecule-a3b1-integrin [49].

Interestingly, Reactome pathway analyses revealed “Integrin cell surface interactions” as over-represented amongst the proteins up-regulated between SCD_PU and SCD. Integrins bind podocyte foot processes to the glomerular basement membrane (GBM) and their impairment leads to detachment of foot processes from the GBM and glomerular filtration problems [52], [53], [54], [55]. By examining the details of the Integrin pathway, we identified PECAM1, SPP1 and BSG among the SCD_PU up-regulated proteins. PECAM1 has been described to directly interact with Integrin αvβ3 to mediate adhesion of leukocytes to the endothelium [56],[57]. Increased levels of PECAM1 is considered a marker for endothelial cells and also for their hyperplasia [58] is in line with SCD hyperplasia associated with hyperfiltration [17] Up-regulated levels of uPA in SCD_PU samples might be further evidence supporting the report that uPA receptor activates integrin αvβ3 in podocytes [59]. Sheinman in his review, described hyperfiltration as a contributing factor that induces endothelial hyperplasia and ultimately glomerular fibrosis [60].

Levels of TFF3 was up-regulated in SCD_PU samples and interestingly increased levels have been associated with CKD [61]. For patients with Type II diabetes which can arise from SCD and mediated by effects on pancreas [62]. It has been hypothesized that the increase in blood glucose and insulin and decrease in short-chain fatty acids may lead to the elevation of serum TFF3 [60].

Urokinase-type plasminogen activator (uPA/PLAU),which is up-regulated in SCD_PU, correlates with disease progression in experimental models of CKD [63]. The level of Its receptor uPAR, decreases in podocyte injury and foot process effacement via activation of α5β3 integrin as reported in a mouse model of lipopolysaccharide (LPS)-induced nephropathy [64], [59].

Comparing the genotypes SS and SC, we identified kidney injury markers Lipocalin-2 (the gene *LCN2* coding the protein NGAL), MMP9, CCN1 and IL1RN (IL1RA) as up-regulated in SS_PU and ANPEP, CLU (Clusterin), IL6, KLK3, RETN and SERPINA3 down-regulated in SS_PU. Furthermore, we could confirm the up-regulation of LCN2 / NGAL in ELISA-based assays. Marouf et al. proposed LCN2/NGAL as a biomarker for nephropathy in sickle cell disease by studying patients with the SS genotype and the HbS/ß0-thalassemia genotype [65] which is also classified as severe and is more prevalent in the Eastern Mediterranean and India [4]. The authors [65] found a correlation of NGAL with microalbuminuria and a better performance of urinary NGAL than serum NGAL for the prediction of vaso-occlusive crisis (VOC) with an Area under the Receiver Operating Characteristic curve for plasma NGAL of 0.69 and for urine NGAL of 0.86. In our experiments, we also observed higher levels of LCN2 in the SC genotype with proteinuria than without in the ELISA assays and additionally higher LCN2 (p=3.42×10^-5^, FDR= 0,0003, ratio=1.42 below our threshold of 1.5) when comparing SCD_PU vs. SCD. NGAL is a transmembrane protein transporting lipophilic substances [65] and has been established as a predictive marker of acute kidney injury [66], [67] and also found to be elevated in SCD during VOC [68]. NGAL is freely filtered through the glomerulus and is re-absorbed by healthy proximal tubular cells and not diseased or injured cells. Lack of re-absorption leading to increased NGAL levels in the urine of patients with kidney injury [66], [65]. By binding to MMP9 (matrix metalloproteinase-9), NGAL inhibits the degradation of MMP9 thus enabling its proteolytic activity and both proteins are correlated with albuminuria in type 1 diabetes patients [69]. Muramatsu and colleagues reported that CCN1 (Cyr61) which is up-regulated in SS_PU, was rapidly induced in rodent proximal tubules and detected in the urine after renal ischemic reperfusion injury [70]. The authors found Cyr61 in the outer rat medulla after ischemic events, in line with medullary VOC in SCD.

In our fluorescence staining and RT-PCR experiments we revealed increased p53 levels in podocytes supplemented with urine from SCD_PU patients. In addition, elevated levels of pro-inflammatory associated mRNAs such as TGFβ in podocytes treated with urine from SCD-PU patients led us to the hypothesis depicted in the scheme in Figure 6: SCD patients with severe inflammation leading to nephropathy is manifested by elevated levels of TGFβ, p53 and also inflammation-associated cytokines RETN, IL1RA, RAGE, TFF3 and PLAU. TGFβ has already been proposed as a urinary marker of renal dysfunction in sickle cell disease [71]. Overstreet et al. proposed that TGFβ contributes to renal injury and fibrosis and that TGFβ activates p53 by phosphorylation which together with SMAD3 transcription factors induce fibrosis in several renal cell types [72]. p53 protein responds to diverse cellular stresses to regulate expression of target genes, thereby inducing cell cycle arrest, apoptosis, senescence and changes in metabolism. Podocyte injury and death has been shown in the mouse by silencing p53 and its inhibitor MDM2 [73]. Further biological processes involved in renal injury include e-organization of the extra-cellular matrix and Integrin cell surface interactions. Kidney injury is manifested by increased levels of LCN2 (NGAL) and decreased levels of CLU. Both our urine SCD, SCD_PU patient and healthy urine analyses coupled with independent analysis of the effect of secreted proteins on human podocytes confirmed the involvement of VEGF (Vascular Endothelial Growth Factor) in the progression of CKD and fibrosis. Interestingly numerous clinical trials are currently exploring the role of VEGF in kidney diseases by using VEGF inhibitors to reduce proteinuria and slow disease progression in conditions like diabetic nephropathy. There is also research into using mesenchymal stem cells over expressing VEGF to promote renal recovery and improve outcomes in chronic kidney disease [74], [75].

**Figure 6:**
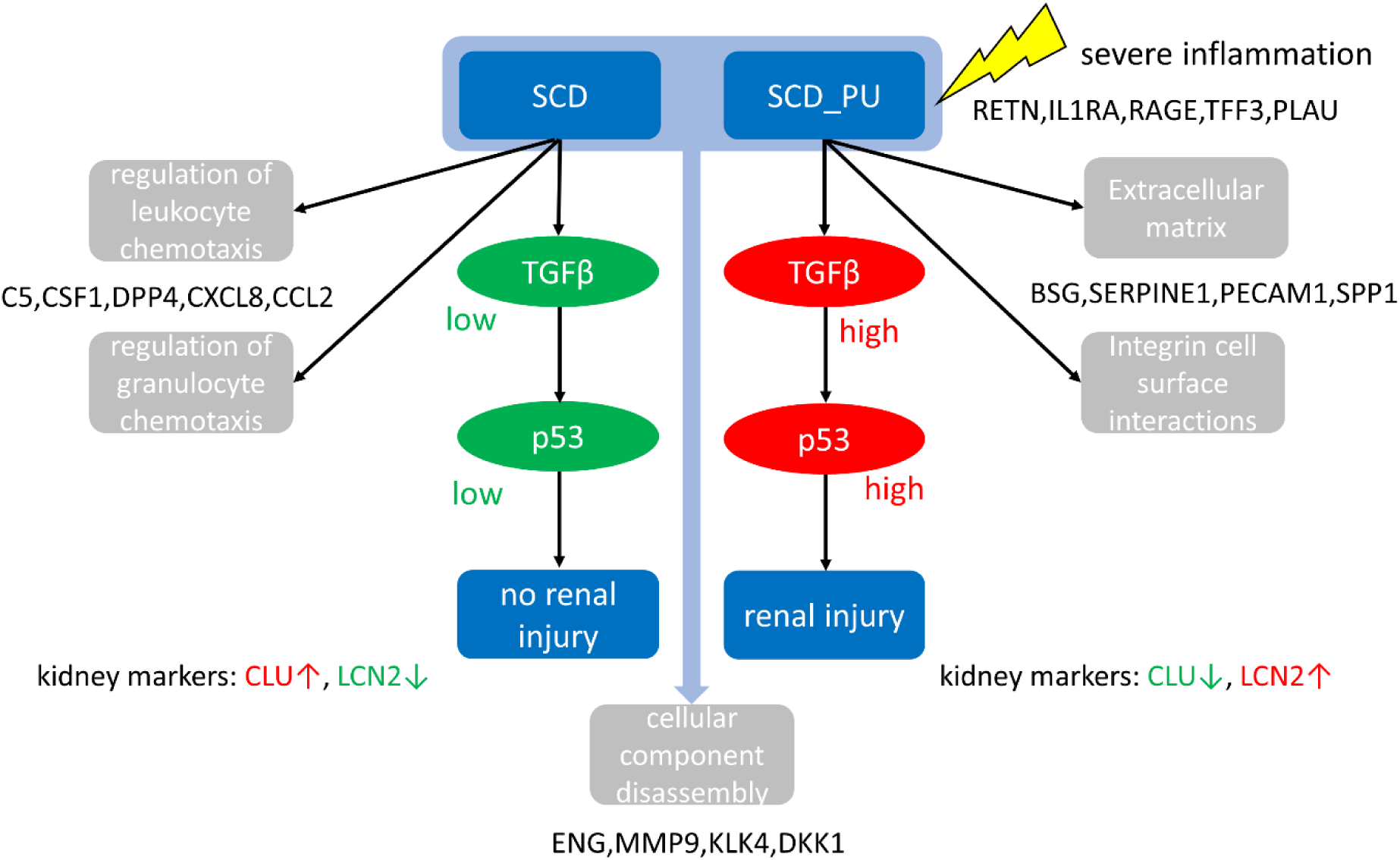
proposed model of how the TGFβ-p53 axis induce renal injury in sickle cell patients. In our cell culture model, we uncovered increased levels of TGFβ and p53 in podocytes supplemented with urine from SCD patients with proteinuria. We therefore hypothesize that SCD patients with more severe inflammation which is manifested by higher levels of TGFβ and other pro-inflammation-associated cytokines RETN, IL1RA, RAGE, TFF3 and PLAU develop nephropathy mediated by p53. Additional biological processes involved in this include organization of the extra-cellular matrix and Integrin cell surface interactions. Kidney injury can be further characterized by high levels of LCN2 (NGAL) and low levels of CLU.

In conclusion, our study revealed distinct urinary kidney injury and cytokine protein profiles between SCD patients without and SCD_PU patients with proteinuria compared to healthy individuals. These profiles also enabled distinguishing SS and SC genotypes. We validated up-regulation of NGAL (encoded by the gene LCN2) in SCD_PU (also in both genotypes SS and SC) and down-regulation of Clusterin with ELISA assays. Follow-up analysis reconstructed a network of urinary secreted proteins in SCD and up-, down or not-regulated in the SCD_PU condition. Proteins up-regulated in SCD_PU were associated with the pathway-“integrin cell surface interactions” which might imply that disrupting the association between hyperfiltration, linking of and integrins to the glomerular basement membrane ultimately leads to glomerular fibrosis.

We finally proposed model of how the TGFβ and p53 axis induce renal injury in sickle cell patients with severe inflammation.

## Author contributions

J.A. and V.B. conceived the study. VB, YD-A, M L, designed and conducted the clinical study, supervised the recruitment and collection of the patient samples.

JA and TK-A stratified patient urine samples.

AIP, CL, RM and CT carried out the experiments and analysed the data. WW provided bioinformatics support. WW, AIP and CT wrote the manuscript.

JA supervised the work, co-wrote the manuscript, edited and gave the final approval.

## Competing interests

The authors declare no competing interests.

## Acknowledgments

James Adjaye acknowledges financial support from the Medical faculty of the Heinrich-Heine University, Duesseldorf.

Vincent Boima acknowledges support from the University of Ghana Medical School and all participants who consented to participate in this study.

## Data availability

The datasets generated during the current study are provided as supplementary data.

## Supplementary Information

**Supplementary Figure 1:**
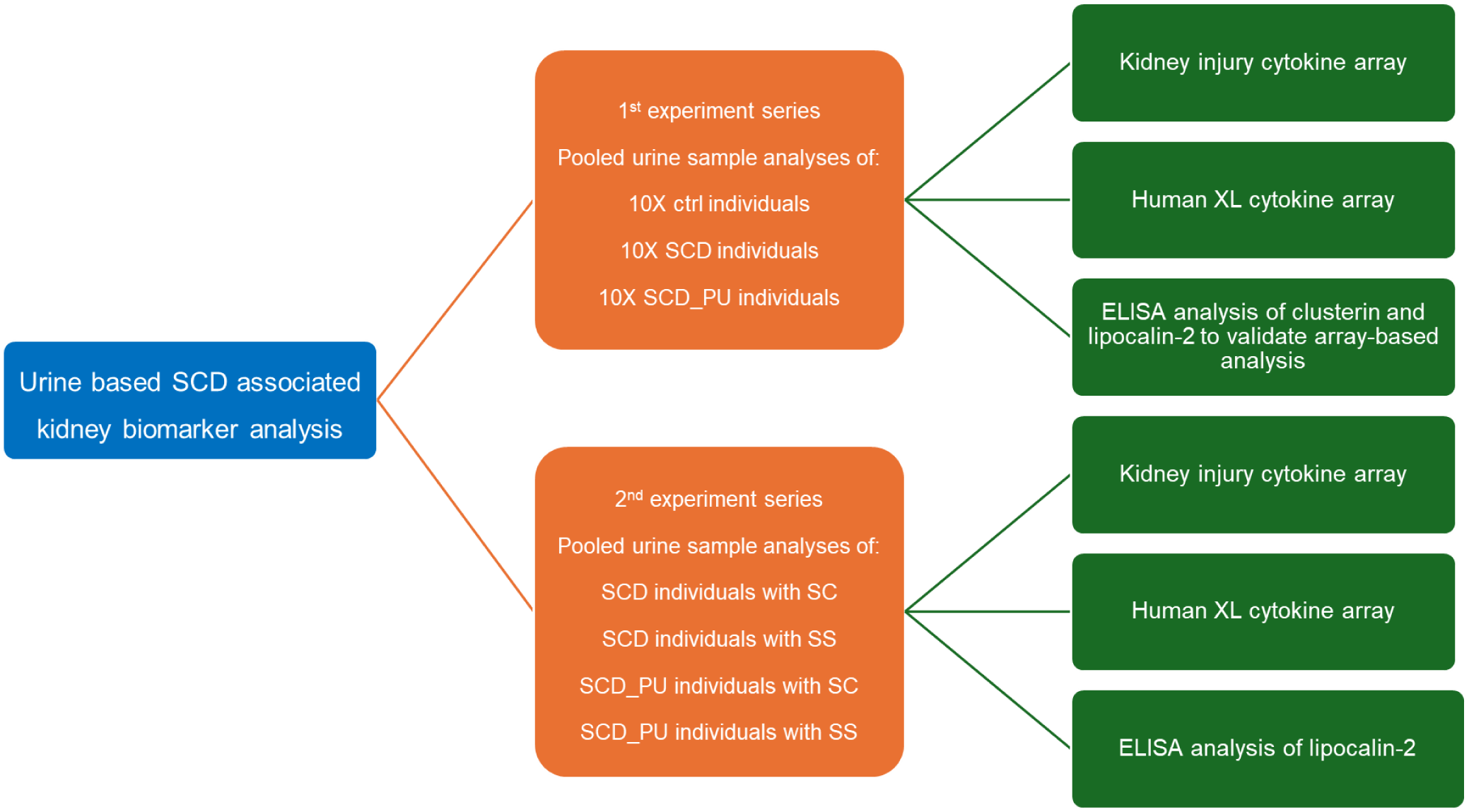
**Flow chart of experiment series of cytokine measurements in urine samples from SCD patients with and without proteinuria and with SC and SS genotypes.**

**Supplementary Figure 2:**
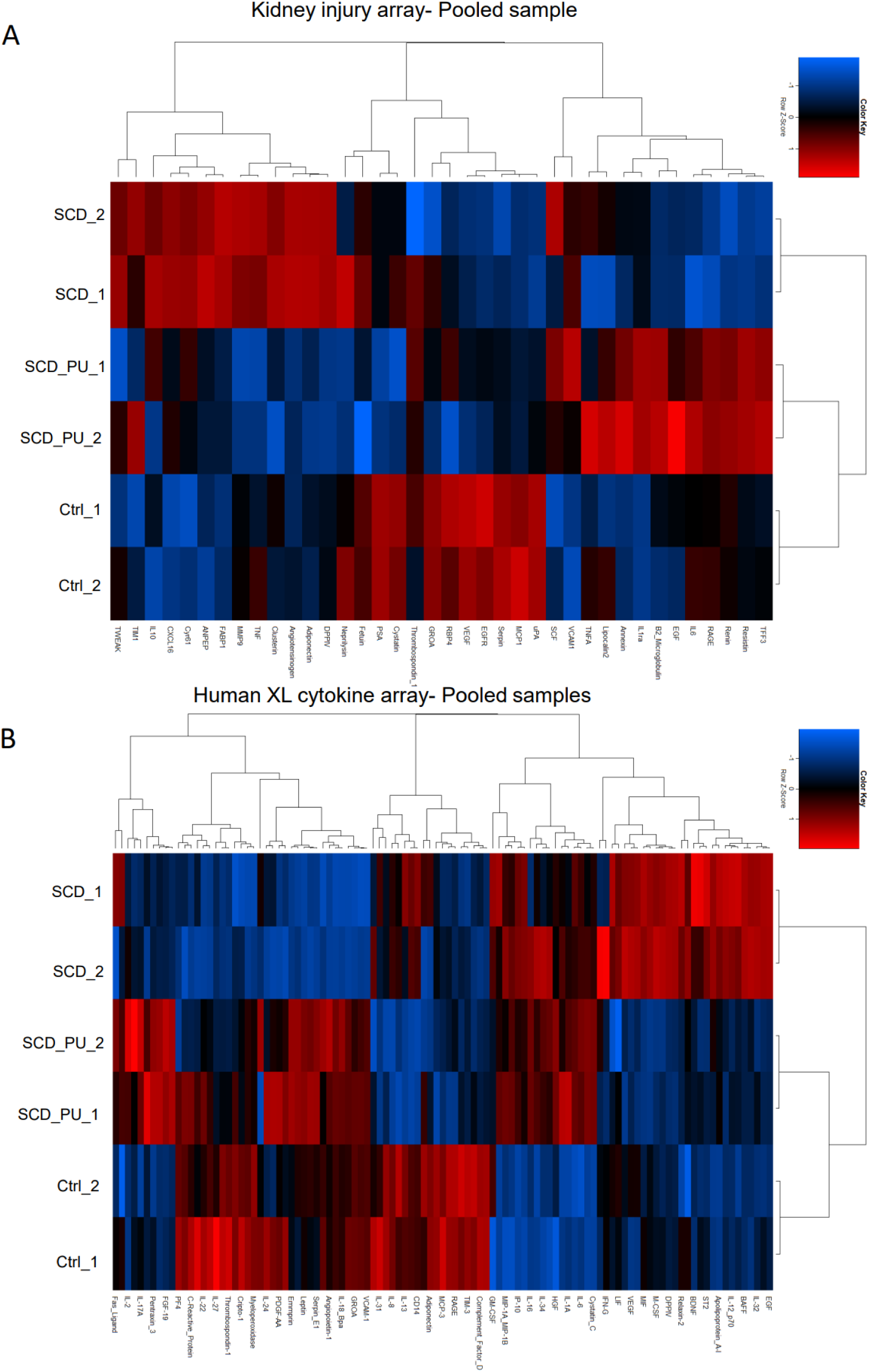
Comparison of protein expression in SCD with and without proteinuria and healthy control. (A) Kidney injury array-Pooled sample. (B) Human XL cytokine array-Pooled samples.

**Supplementary Figure 3.**
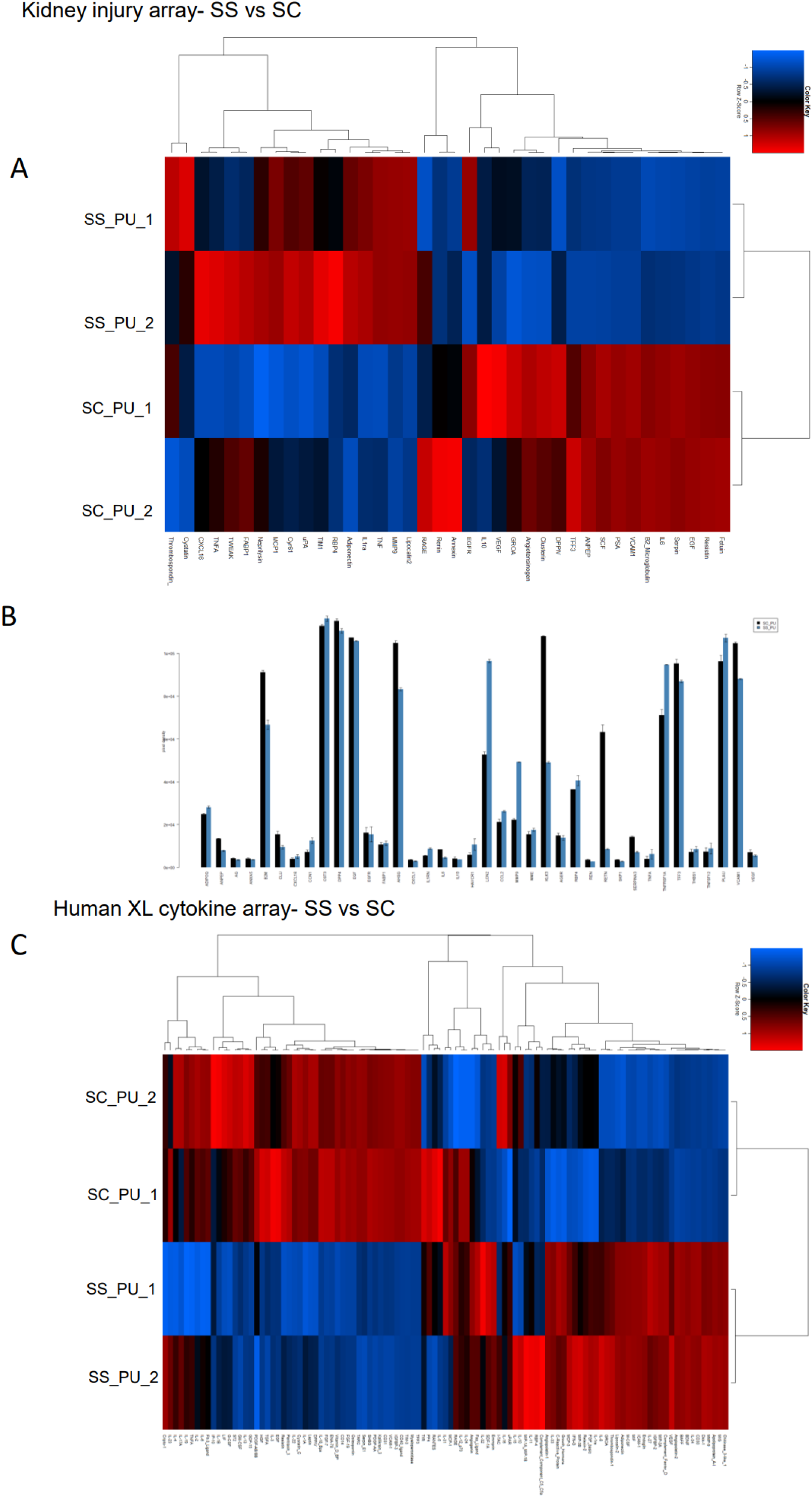
(linked to main. **fig 1+2): Comparison of protein in genotypes SS and SC.** (A) Heatmap depicting expression of proteins on the Kidney injury array-SS vs SC. (B). Bar chart depicting expression of proteins on the Kidney injury array-SS vs SC. (C) Heatmap depicting expression of proteins on the Human XL cytokine array-SS vs SC.

**Supplementary Figure 4.** (figS4.pdf, linked to main. **fig 1, 2 and 4)**: Scanned cytokine arrays of experiments in this study:(A) Pooled urine samples of SCD_PU, SCD and Ctrl on kidney protein array. (B) Pooled urine samples of SCD_PU, SCD and Ctrl on Human XL cytokine array. (C) Pooled urine samples of genotypes SS and SC on kidney protein array. (D) Pooled urine samples of genotypes SS and SC on Human XL cytokine array.

## Supplementary Tables

**Supplementary Table 1 (table S1.xlsx): Antibodies and primers.**

**Supplementary Table 2 (tableS2_kmembrane_SCD_PU_SCD_ctrl.xlsx): mean values and statistical test results of (a) SCD_PU vs. control, (b) SCD vs. control and (c) SCD_PU vs. SCD on kidney injury marker array.**

**Supplementary Table 3 (tableS3_humanXL_SCD_PU_SCD_ctrl.xlsx): mean values and statistical test results of (a) SCD_PU vs. control, (b) SCD vs. control and (c) SCD_PU vs. SCD on human XL cytokine array. Corresponding metascape enrichment results are in sheets (d-f)**

**Supplementary Table 4 (tableS4_SS_PU_vs_SC_PU.xlsx): mean values and statistical test results of genotype SS and SC comparisons (a) SS_PU vs. SC_PU on kidney injury marker array, (b) SS_PU vs. SC_PU on human XL cytokine array.**

## Notes

### Competing Interest Statement

The authors have declared no competing interest.

