## Supplementary Figure 4 for "Urinary proteins from Sickle Cell patients induce inflammation and kidney injury via the TGFβ-p53 axis in a podocyte cell culture model"

A

Kidney Biomarker Array :

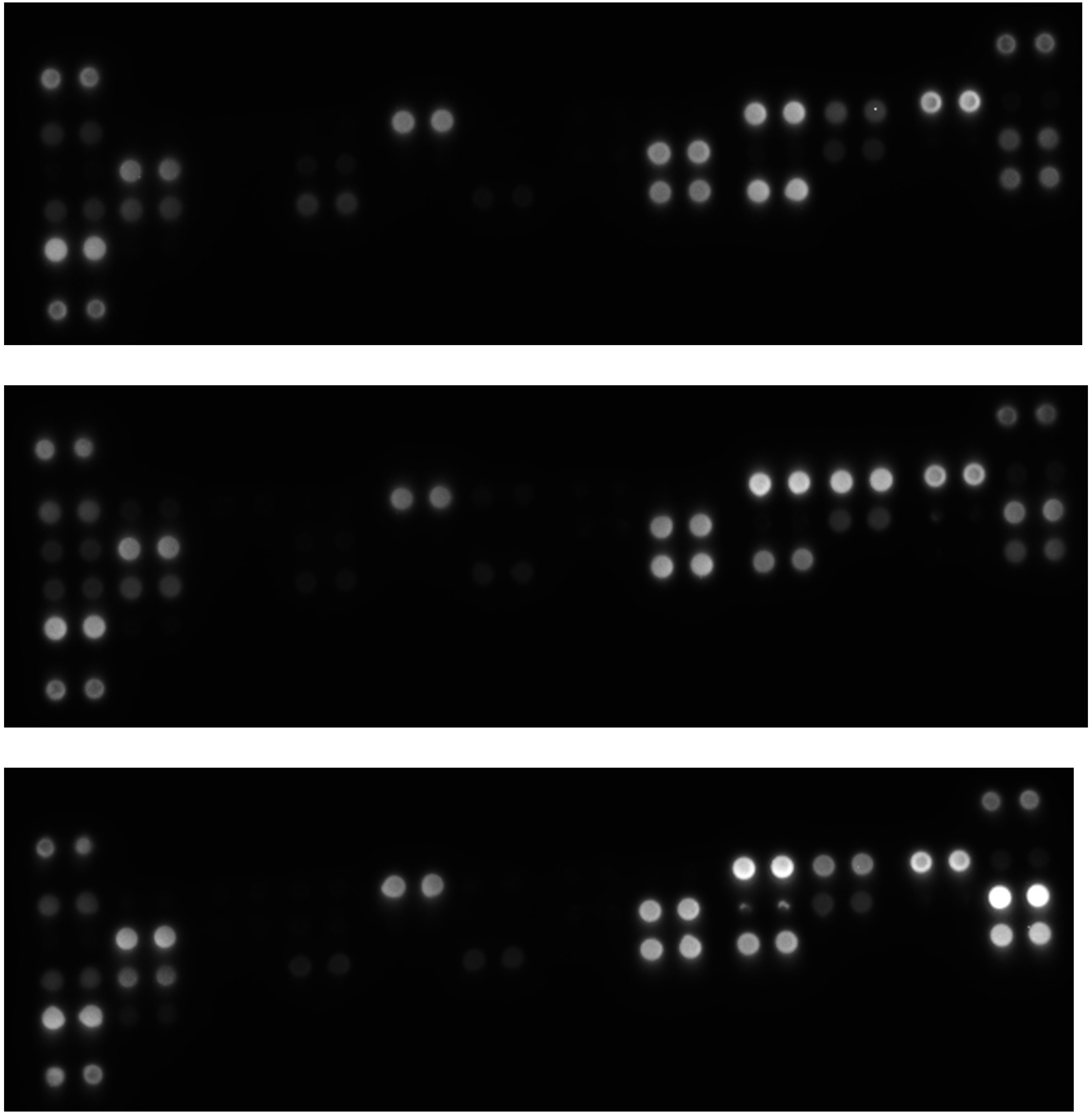

SCD_PU

SCD

Ctrl

B

Human XL cytokine array :

SCD_PU

SCD

Ctrl

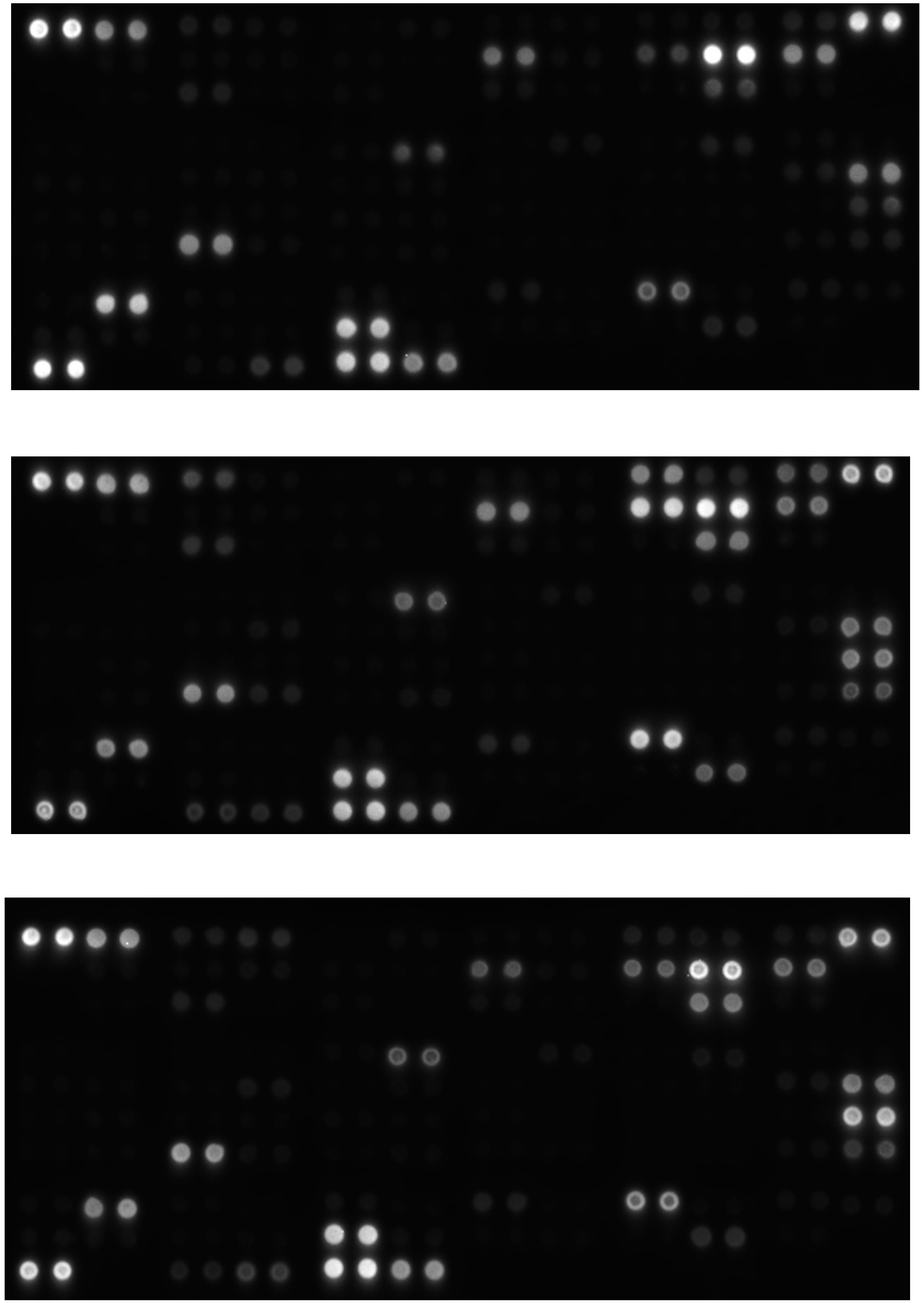

C

Kidney Biomarker Array :

SS_PU

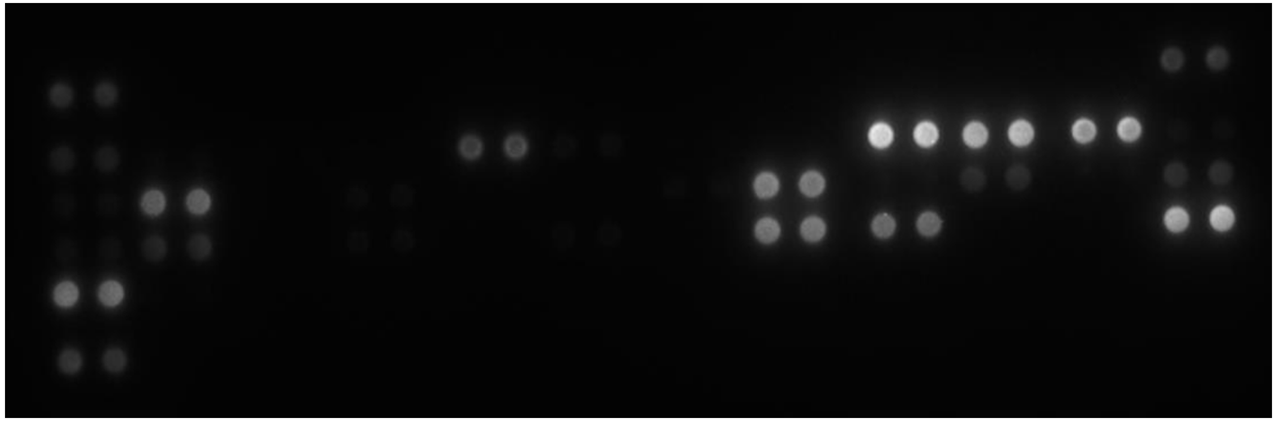

SC_PU

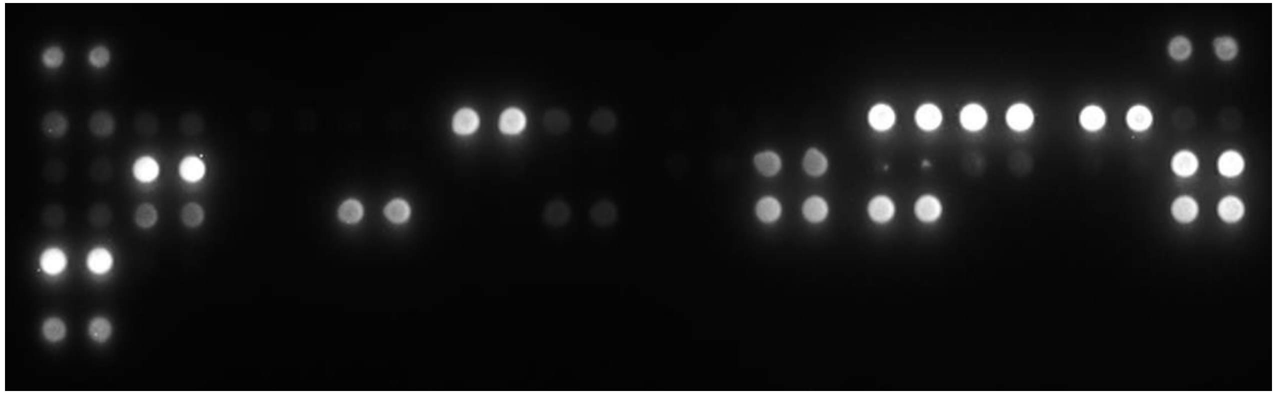

D

Human XL cytokine array :

SS_PU

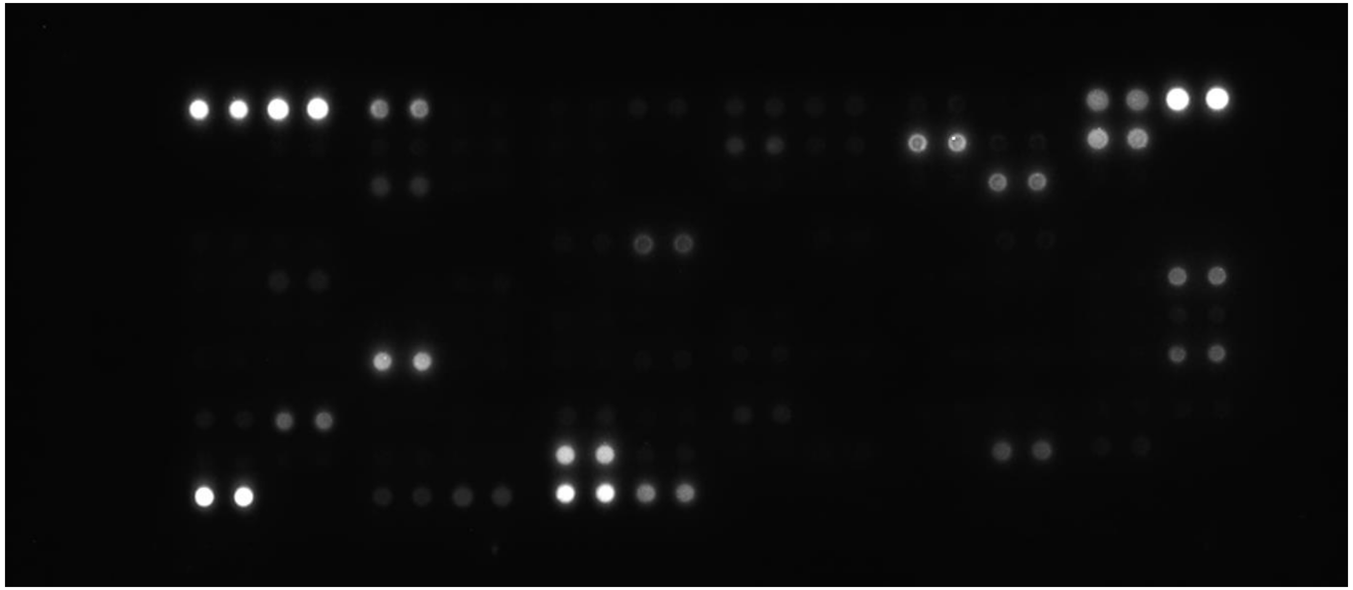

SC_PU

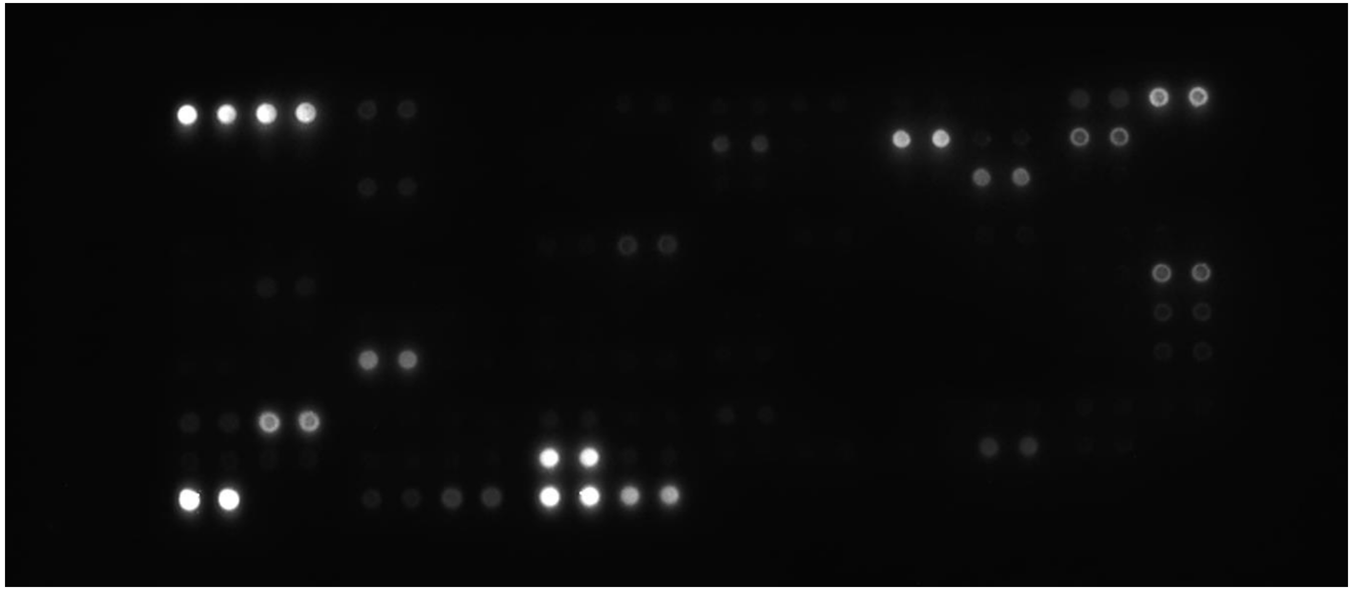
